# A cascade of attentional and default mode network dynamics following the resolution of perceptual ambiguity

**DOI:** 10.64898/2026.09.26.754627

**Authors:** Adam Ponzi, Katsunori Miyahara, Keisuke Suzuki

## Abstract

The neural mechanisms underlying spontaneous perceptual switches during bistable perception, particularly the dynamic interplay between large-scale brain networks, remain incompletely understood. Here, using functional magnetic resonance imaging (fMRI) in six participants and a bistable kinetic depth effect (KDE) stimulus, we investigated the spatiotemporal dynamics of brain activity associated with spontaneous perceptual reversals compared to physically induced (replayed) switches. Leveraging a fine-grained functional brain parcellation (MD758) and time-resolved analysis including noise correlations, we identified a distributed network encompassing occipital (MT/MST, LOC), parietal (IPS), and frontal (FEF, IFC, DLPFC) regions showing differential activity during spontaneous versus replayed switches. This network topography qualitatively overlapped a prior review-derived synthesis of perceptual-switching regions. Temporal decomposition of the differential fMRI signal further revealed a characteristic network cascade after the perceptual report: positive dorsal attention (DAN) and visual (VIS) network contrasts peaked approximately 4 s after the report alongside a negative default mode network (DMN) contrast; a slower, more modest ventral attention (VAN)–frontoparietal (FPN) component was observed around 10 s post-report but was weak and is treated as preliminary. The early 4 s profile was recovered across raw-signal, PCA-denoised, and noise-correlation analyses—three related, not fully independent analyses—with the noise-correlation analysis indicating that it is not fully attributable to shared motor-report signals. These findings delineate the temporal sequence of network recruitment following spontaneous perceptual transitions and suggest that interplay between attentional control and default mode networks accompanies the resolution of perceptual ambiguity.

**Highlights:**

- Distributed cortical cascade of network engagement follows spontaneous bistable KDE perceptual switching.
- Positive DAN–VIS and negative DMN network contrasts peak 4 s post-report.
- A preliminary, slower VAN–FPN component is observed around 10 s post-report.
- Noise-correlation analysis indicates the early network profile is not fully explained by shared motor-report signals.
- Findings temporally dissect the review-derived BRAS network of bistable perception.

## 1 Introduction

When humans view ambiguous stimuli such as the Necker cube, perception spontaneously alternates between competing interpretations even when the physical stimulus remains unchanged [1–3]. Standard neural models attribute these alternations to mutually inhibitory populations encoding each percept, with switches arising from noise, adaptation, or other perturbations that destabilise the currently dominant attractor state [4–8]. While such mechanistic accounts capture the local dynamics of competition, the precise spatiotemporal choreography by which large-scale brain networks coordinate around these endogenous perceptual transitions remains elusive. Understanding how distinct brain systems coordinate over time to resolve perceptual ambiguity is critical for elucidating the neural basis of conscious perception and cognitive control.

A long-standing debate concerns the locus of these switches. One view emphasises early sensory processing stages, where competition between feature- or eye-selective representations may be sufficient to drive alternations [9–13, 33]. The competing view holds that switches are driven by top-down control from frontoparietal regions implicated in attention, working memory, and decision-making [14–18, 20, 21]. Although evidence supports top-down modulation of visual cortex activity [22, 23] and the suppression of incongruous percepts at higher cognitive levels [19, 24–26], a critical gap remains in understanding the *dynamic* network interactions underlying these processes. Moving beyond static loci of activity requires elucidating the temporal sequence of network engagement during the moments preceding and accompanying a spontaneous perceptual switch.

Previous studies comparing spontaneous and replayed switches have consistently identified a right-lateralised frontoparietal network—including the inferior frontal cortex (IFC), frontal eye field (FEF), dorsolateral prefrontal cortex (DLPFC), and intraparietal sulcus (IPS)—as differentially engaged during endogenous transitions [15, 34, 35, 37]. A review and synthesis by Brascamp and colleagues [37] consolidated these findings into a set of regions, hereafter referred to as the BRAS network, that are commonly reported during spontaneous versus replayed switching across stimulus paradigms. These regions overlap substantially with the dorsal (DAN) and ventral attention networks (VAN) [39, 40], linking multistable perception to large-scale attentional control systems [2, 15, 41– 43], with right parietal regions particularly implicated in perceptual awareness [25, 44, 45]. Complementing this picture, coupling of the default mode network (DMN) with lateral occipital and frontoparietal networks has been linked to bistable perception [81], and the DMN is typically suppressed during externally directed, attentionally demanding tasks [46, 78], an effect often interpreted in the literature as reflecting an interruption of internally directed thought processes, though this specific cognitive interpretation is inferred from the DMN’s role in other tasks rather than directly tested. However, how these distinct networks—DAN, VAN, DMN, and the sensory regions they modulate—interact dynamically over time during a spontaneous switch remains a key unanswered question.

A further methodological challenge complicates this picture. Because perceptual switches are typically identified from an explicit behavioural report, any report-locked neural signal risks conflating processes tied to the perceptual transition itself with those tied to introspection, decision-making, and motor preparation for the report—a concern raised most directly by no-report paradigms, which have shown that frontal activity linked to reporting a rivalry switch can be dissociated from, and mistaken for, the perceptual switch itself, whereas occipital and parietal activity persists without report [50]. Any design relying on an explicit report, including the present one, must confront this issue directly rather than treat it as a peripheral caveat.

To address this gap, we investigated the network dynamics of spontaneous perceptual switching using a kinetic depth effect (KDE) stimulus, a structure-from-motion display that elicits bistable rotational percepts [27–32, 36]; because KDE is a motion-based paradigm distinct from the binocular-rivalry and ambiguous-figure displays that dominate the bistable-perception literature, it additionally offers a test of whether the frontoparietal network implicated in that literature generalises beyond the paradigms used to define it. We compared brain activity during spontaneous reversals with activity during externally induced, replayed reversals in a control stimulus [15, 37]. Because the replayed control used asymmetric rather than symmetric rings (see Methods), this contrast does not fully separate switch origin from low-level stimulus geometry. We combined functional magnetic resonance imaging (fMRI) with a fine-grained functional parcellation derived from resting-state connectivity (MD758 [84]) and time-resolved analyses including noise correlations and event-related averaging across distinct network nodes; together with the replayed-switch control, the noise-correlation analysis is intended to address the report-locking concern raised above, since it removes the trial-averaged, time-locked component of activity shared by both conditions (see Methods). By relating our findings to the review-derived BRAS regions and to canonical resting-state networks, we aimed to map the sequence and functional coupling dynamics of large-scale networks that accompany endogenous perceptual transitions.

## 2 Results

Before presenting the results, we briefly describe the two perceptual tasks used in this study. In both tasks, participants reported when they perceived a switch in the rotational direction of a visual stimulus. In the *spontaneous* task, the rotation was fully ambiguous, so that perceptual switches arose endogenously from intrinsic neural dynamics. In the *replayed* task, the stimulus rotated in a defined direction that was reversed at pre-programmed intervals matched in mean to the spontaneous reversals, requiring participants to report externally induced perceptual transitions (see Methods).

### 2.1 Perceptual Switch-Related Brain Activity

To characterise how brain activity changes during perceptual switching, we analysed whole-brain signal fluctuations time-locked to reported transitions. We first examined overall activation patterns using peri-switch time histograms (PSTHs). We then applied principal component analysis (PCA) to extract dominant report-locked response patterns, and finally examined time-resolved functional connectivity (FC) to assess how between-area interactions evolved around the perceptual transition.

#### Whole-brain PSTHs reveal stronger and earlier modulations under spontaneous switching

We computed report-locked PSTHs of standardised fMRI activity for all 758 MD758 regions, then averaged into the 90 AAL regions for visualisation (Fig.1A,B). Both spontaneous and replayed switching produced strong report-locked modulations across multiple regions, but these modulations were markedly stronger and earlier in the spontaneous condition. Peak modulations occurred approximately 4 s after the perceptual report, consistent with the haemodynamic response delay. In addition, the spontaneous task—but not the replayed task—exhibited a small reduction in area-averaged activity in the few seconds *preceding* the report (Fig.1C), consistent with an anticipatory preparatory state that is absent when transitions are externally driven; because spontaneous trials are also self-timed and temporally unpredictable while replayed trials are externally timed, this prereport reduction cannot be fully attributed to perceptual processes as opposed to motor-timing or temporal-expectation differences between the two tasks (see Discussion). The trial-by-trial standard deviation of activity (Fig.1D,F) was also more strongly and more rapidly suppressed around 4 s post-report in spontaneous switching, indicating that the post-switch dynamical evolution was highly stereotyped across trials.

**Figure 1.**
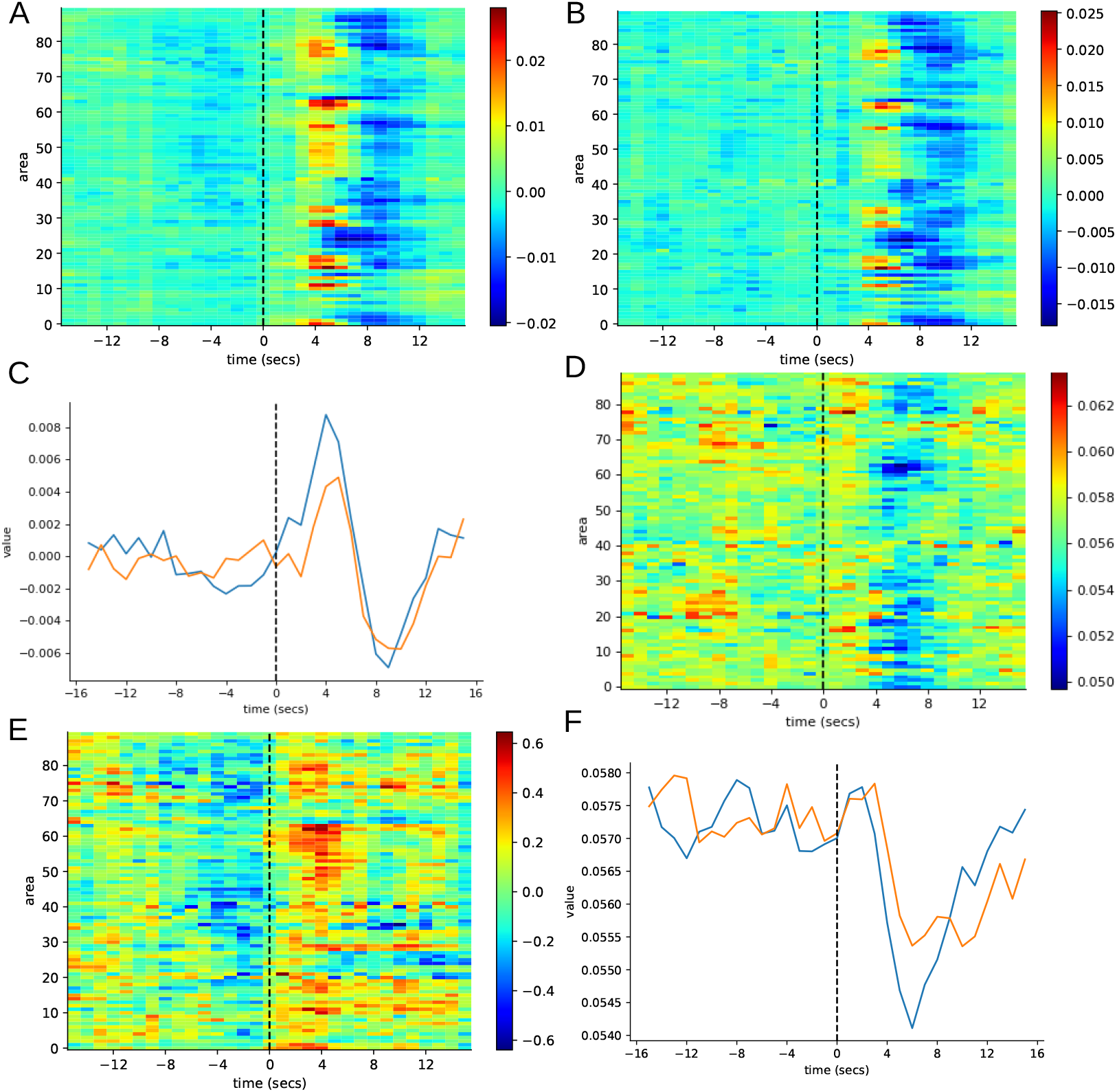
Overall brain activity. (A,B) Cross-trial PSTH of standardised brain activity locked to perceptual report at time *τ* = 0 in (A) spontaneous and (B) replayed perceptual switching tasks. (C) Mean of the cross-trial brain-activity PSTH averaged across brain areas in spontaneous (blue) and replayed (orange) perceptual switching. (D) Standard deviation of brain activity locked to perceptual report at time *τ* = 0 in spontaneous perceptual switching. (E) Task *differential z-scores*, denoted dSig, calculated from the standardised brain activity shown in (A,B). (F) Mean of the standard deviation of brain activity averaged across brain areas in spontaneous (blue) and replayed (orange) perceptual switching. (A–F) Results are calculated for individual subjects separately then averaged. All calculations use 758 brain areas determined by MD758 parcellation of fMRI voxels, averaged down to 90 brain areas of the AAL parcellation for visualisation purposes.

#### A distributed differential network distinguishes spontaneous from replayed switching

To isolate brain regions that were differentially engaged between the two conditions, we computed a task-differential signal (dSig) by subtracting the replayed PSTH from the spontaneous PSTH at each time point. Statistical significance was assessed against a null distribution of 3000 surrogate datasets generated by randomly resampling trials, with replacement, from the pooled trials of both conditions (see Methods). The resulting z-score map (Fig.1E) showed areas with *greater* activity in the spontaneous condition 3–4 s after the report. Greater activity in the replayed condition was weaker and less consistent in time; in the combined-subject test (Fig.2A), only a few such areas reached significance before the report, no more than expected by chance (see Methods).

**Figure 2.**
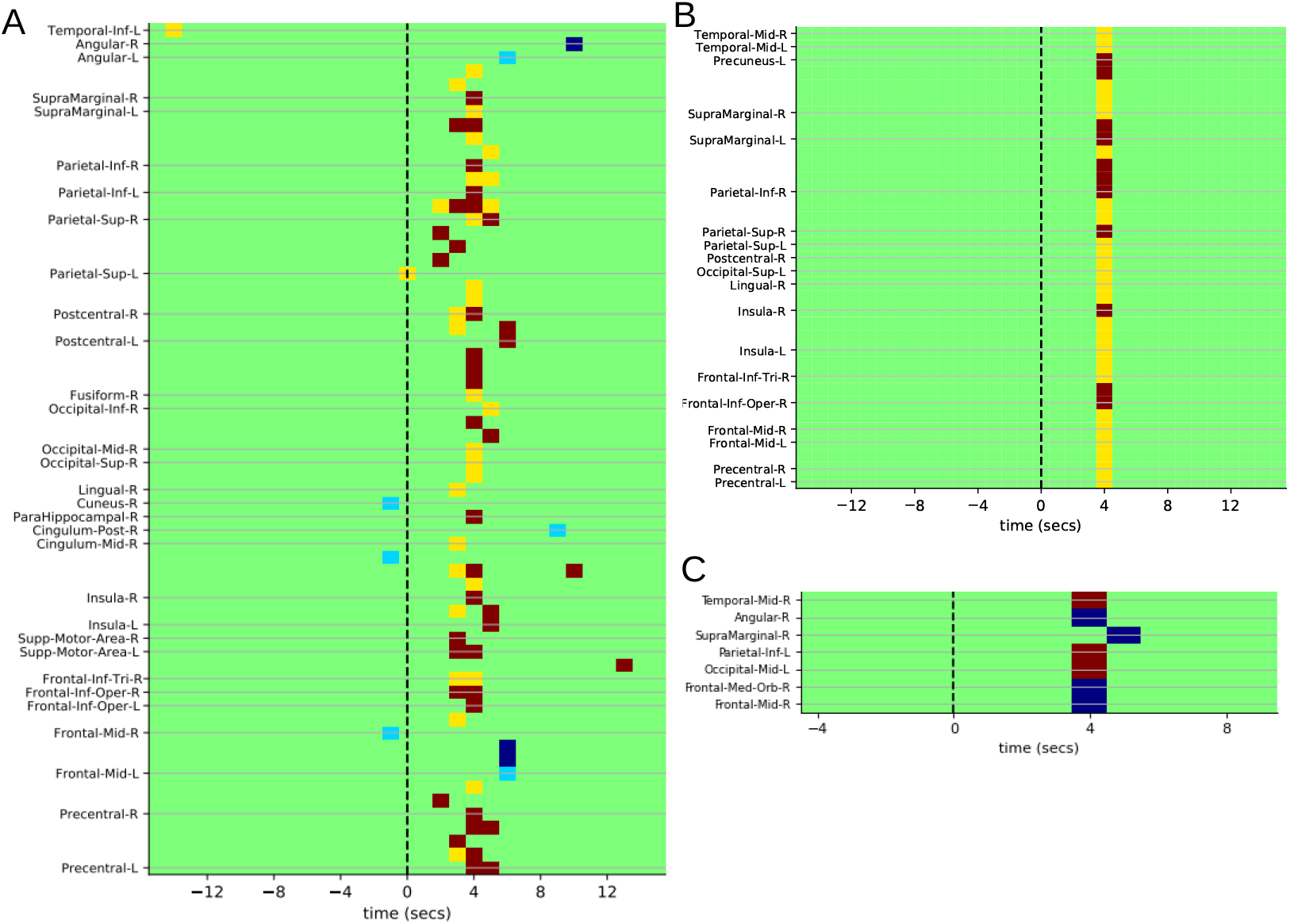
Significantly positively and negatively *differentially* activated brain areas in spontaneous compared to replayed perceptual switching locked to perceptual report at *τ* = 0, calculated from (A) the raw signal (dSig), (B) the eigenvalue weighted sum of the first five PCs (dPC1-5), (C) the first FC component (dFC1). (A,B) Comparison with *N*_sur_ = 3000 surrogates obtained by resampling trials, with replacement, from the pooled spontaneous and replayed trials. (C) Comparison with *N*_sur_ = 500 surrogates. (A,B) In areas shown red (yellow) the actual dPSTH was more positive than *N*_sur_ (*N*_sur_ − 1) surrogates; in areas shown blue (cyan) it was more negative than *N*_sur_ (*N*_sur_ − 1) surrogates. (C) Only the most extreme level is shown: red (blue) areas were more positive (negative) than all *N*_sur_ surrogates. All MD758 areas with at least one significant point in the ± 15 s window around report are included. Labels are from the AAL parcellation; each AAL region contains multiple MD758 areas. Labelled brain areas continue above each labelled horizontal line until the next line. Results are for combined subjects: individual subject dPSTH are averaged and the average compared with the same calculation made from surrogates (see Methods).

#### Cortical mapping of the dSig differential network at the 3-s peak

Examining all 758 MD758 regions in the dSig analysis (Fig.2A) confirmed a broad network spanning occipital, parietal, and frontal areas that was more active during spontaneous than replayed switching, peaking 3–4 s after report. Only a small subset of regions (e.g., a left mid-frontal area at ~6 s post-report) showed the opposite pattern. Mapping the most significantly activated regions at *τ* = 3 s onto the cortical surface using the HCP 360-area parcellation [38, 60] (Fig.S1) revealed FEF, SCEF, LIPv, 7PC, PFt, PFop, IFG areas 44 and 45, and frontal opercular areas FOP3–5—regions commonly implicated in bistable perception (see Discussion).

#### Principal components isolate a sharper, lower-noise differential network

The coactivation of many regions in the raw dSig analysis suggested that principal component analysis (PCA) could compress the report-locked response into a small number of dominant patterns. We computed PCA on the time-series cross-correlation matrix across all 758 MD758 regions in the spontaneous condition, then projected both conditions onto the resulting components. The first principal component (PC1) exhibited a global, triphasic time course: suppression before the switch, a sharp peak at 4 s post-report, and a return below baseline near 10 s (Fig.3A). PC2 showed more sustained modulations between 4 and 10 s post-report, with hemispheric asymmetry (Fig.3C), possibly reflecting longer-lasting reconfiguration of cognitive control circuitry following the transition.

**Figure 3.**
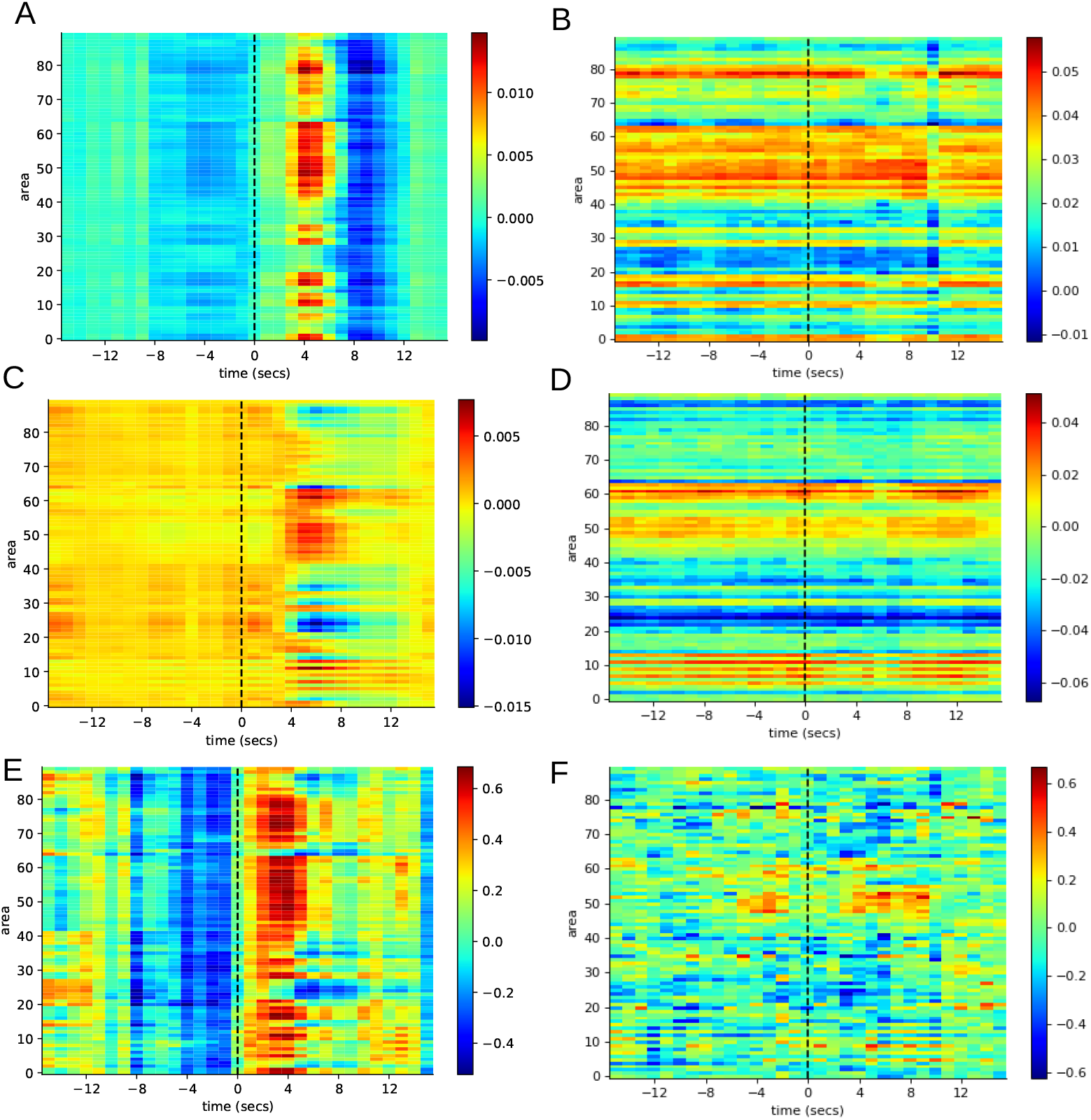
Principal component PSTHs. (A,C) Cross-trial PSTH of the temporal projection of the first (PC1, A) and second (PC2, C) principal components of brain activity in spontaneous perceptual switching. (B,D) First (FC1, B) and second (FC2, D) functional connectivity (cross-trial noise correlation matrix) principal components at each time, in spontaneous switching. (A–D) Subjectaveraged. (E,F) Task *differential z-scores* from (E) the eigenvalue-weighted sum of the first five PCs (dPC1-5) using 3000 surrogates and (F) the first noise-correlation principal component dFC1 using 500 surrogates. Here z-scores are calculated per subject before averaging. All calculations use MD758 parcellation; averaged down to 90 AAL regions for visualisation.

#### The dPC1-5 differential at the 4-s peak

Combining the first five PCs in an eigenvalue-weighted sum produced a denoised differential signal, dPC1-5, evaluated for significance against 3000 surrogates (Fig.3E). Although fewer regions reached significance than in dSig, those that did were more tightly clustered in time around the 4 s peak and in space, consistent with an improved signal-to-noise ratio (Fig.2B). The significant regions at *τ* = 4 s (Fig.4A) included bilateral MT and MST, FEF, IPS, AIP, and inferior frontal regions—consistent with previous literature on bistable perception and pointing to an interplay between bottom-up sensory and top-down attentional control processes (see Discussion).

**Figure 4.**
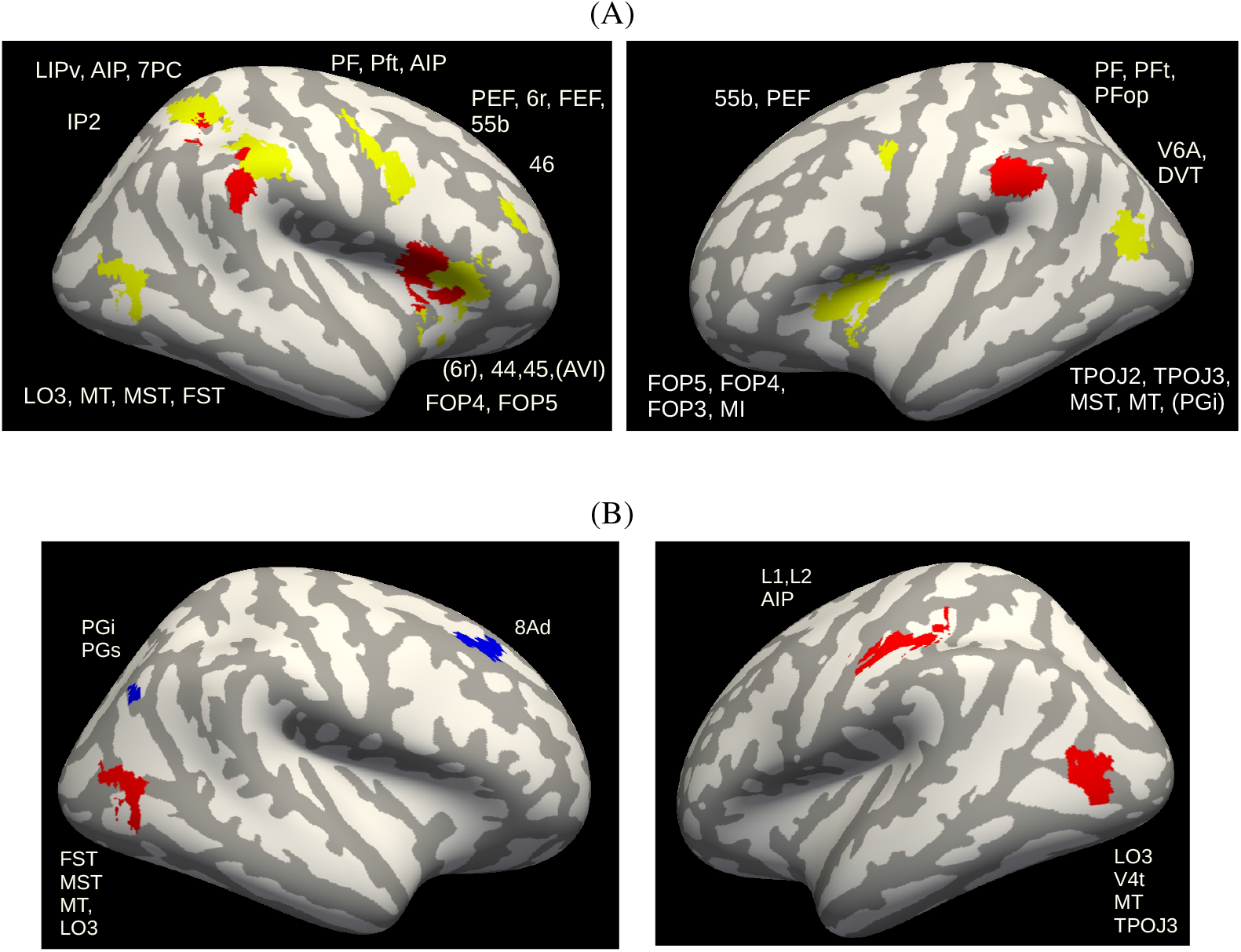
Cortical localisation of the task-differential network at the *τ* = 4 s post-report peak, labelled according to the HCP 360-area parcellation [38, 60] (all calculations use the MD758 parcellation). **(A)** Areas significantly differentially activated in spontaneous compared to replayed perceptual switching using dPC1-5 (eigenvalue-weighted sum of the first five PCs), corresponding to Fig.2(B); red (yellow) areas: actual dPSTH more positive than 3000 (2999) surrogates. **(B)** The same contrast computed from the first noise-correlation principal component, dFC1, corresponding to Fig.2(C); red (blue) areas: actual dPSTH more positive (negative) than 500 surrogates.

#### Noise-correlation connectivity reduces the contribution of motor-related modulations

The principal components derived from the full-task correlation matrix may capture substantial variance from the motor response associated with the button press. Because this response is largely shared between spontaneous and replayed conditions, it may not be fully removed by subtracting replayed from spontaneous PSTHs—particularly if the temporal dynamics of motor preparation differ subtly between conditions. To reduce the contribution of this shared motor component, we instead computed report-locked *noise correlation* matrices: at each time point relative to the report, we subtracted the trial-averaged PSTH from each individual trial and correlated the resulting residuals across all area pairs (Fig.5A). This yields a time-resolved sequence of connectivity matrices that, by construction, removes the trial-averaged (mean) motor-report component shared across trials of a condition; residual trial-to-trial motor variability, or condition-level differences in motor-timing predictability, could in principle remain (see Discussion). We then applied PCA to each matrix independently, producing time-specific functional connectivity components (FC1, FC2, and so on). Because each FC component is an eigenvector, its task-differential (dFC1) measures a difference in how strongly each area loads on the leading mode of trial-to-trial covariation, not a difference in activity; below, ‘more strongly modulated’, ‘positive’, and ‘negative’ refer to this loading difference.

**Figure 5.**
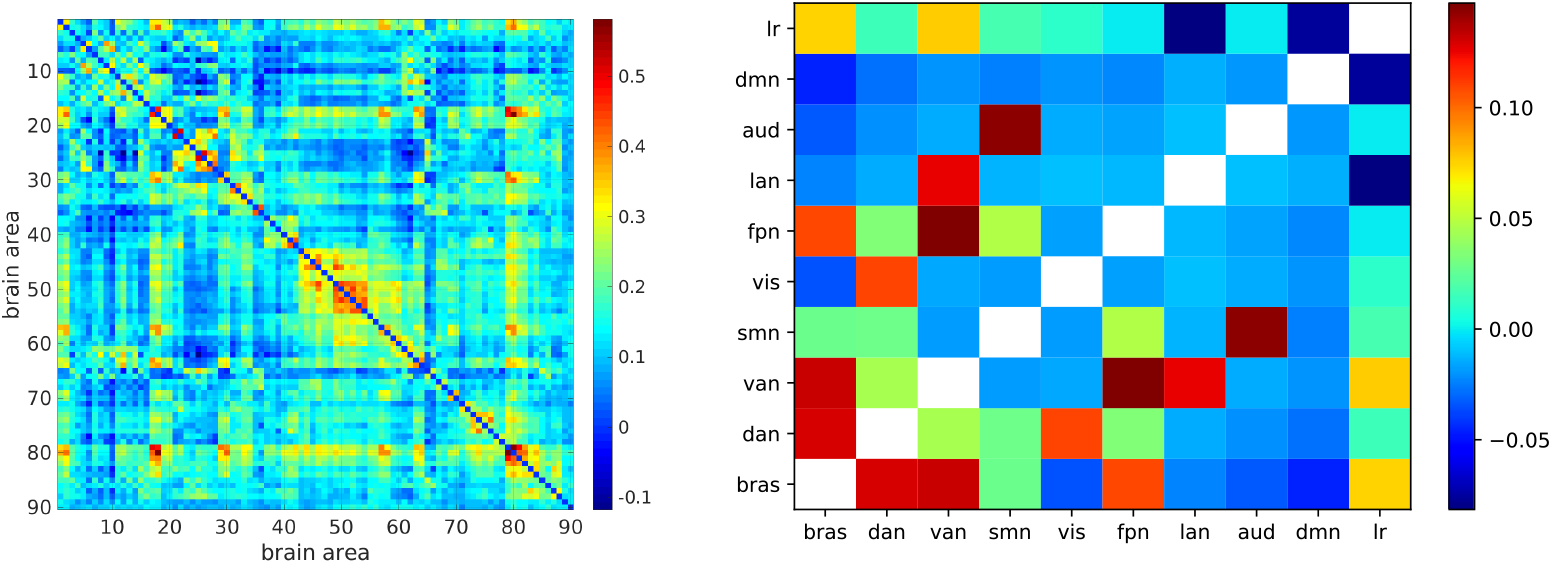
(A) Example noise correlation matrix calculated from MD758 parcellation, averaged to 90 AAL areas for visualisation, for replayed perceptual switching at *τ* = 0 (subject average). (B) Structural correlation coefficients between the BRAS network and the RSNs. The correlation between these networks and a left–right (LR) lateralisation vector is also shown: positive when a network is more situated in the right hemisphere. Networks are binary vectors over the brain areas, normalised so that brain areas in neither of a pair contribute positively to their correlation. Unity diagonal removed.

#### Noise-correlation components identify bilateral MT as central to the cascade

As expected, FC1 and FC2 (Fig.3B,D) showed much weaker temporal modulation than PC1 and PC2, consistent with the removal of the trial-averaged response. The two decompositions are nevertheless computed from covariation among the same areas, so FC1 and PC1 are not fully independent measures of the underlying signal; FC1’s distinct contribution is specifically to remove the trial-averaged motor component that PC1 retains, rather than to constitute a wholly separate replication (see Discussion). The task-differential of FC1 (dFC1, evaluated against 500 surrogates; Fig.3F) identified parietal and occipital areas as more strongly modulated under spontaneous switching, while paracentral lobule and temporal pole regions were reduced. Mapping these onto the cortical surface at *τ* = 4 s (Fig.4B) revealed that bilateral MT was the most strongly modulated region, consistent with its established role in motion and structure-from-motion processing. The map also included a left anterior parietal cluster around AIP, close to sensorimotor cortex (Fig.4B), so a residual report-related sensorimotor contribution to dFC1 cannot be excluded.

#### Convergence across analyses: a coherent occipital–parietal–frontal network

Across the three analyses—the raw differential signal (dSig), the PCA-denoised differential (dPC1-5), and the noise-correlation differential (dFC1)—a convergent picture emerges, though, as noted above, these are complementary views of the same underlying signal rather than fully independent replications. Spontaneous perceptual switching engages a broad cortical network peaking around 4 s post-report. Motion-selective occipital areas (MT, MST, LO3) and AIP were present in both the dPC1-5 and dFC1 maps at 4 s (Fig.4), and occipital areas were also significant in dSig (Fig.2A); further parietal regions (7PC, PF) and inferior frontal and premotor regions (IFC, FEF) appeared in dSig and dPC1-5 but not in dFC1, and SCEF appeared only in dSig. The noise-correlation approach—which removes the trial-averaged motor component by construction—indicates that at least this occipital and intraparietal core is not fully explained by the report response itself, though it does not rule out contributions from trial-to-trial motor variability.

### 2.2 Cross-subject consistency supports the robustness of the cascade

The above analyses combine three complementary measures—dSig, dPC1-5, and dFC1—that differ in how they handle shared time-locked signals and noise. To identify which yields the most robust read-out of perceptual switch-related dynamics, we computed an inter-subject consistency (ISC) measure (Fig.6). For each time point relative to the perceptual report, we computed the pairwise spatial-similarity score defined in Methods from differential PSTH z-scores across brain areas, then averaged across all 15 subject pairs. High ISC at a given time descriptively indicates that subjects enter similar brain states at that moment.

The dSig measure (green) showed a clear ISC peak around 4 s post-report. dPC1 alone (blue) reached about half the dSig peak but rose earlier, possibly capturing earlier components of the response. The eigenvalue-weighted sum dPC1-5 (orange) rose rapidly after the report and achieved the highest overall consistency, indicating that combining multiple PCs effectively distils task-relevant patterns shared across individuals while suppressing noise. However, to the extent that the motor response differs between conditions (see above), these consistency measures may be dominated by consistency in the report-locked motor response across subjects. dFC1 (red)—which removes the trial-averaged, report-locked component—showed only a small rise after the report (peak ≈ 0.09), comparable to fluctuations at other time points, and therefore provides at most weak evidence for a shared spatial pattern across subjects once that component is removed. Because no inferential null distribution or confidence interval was computed for ISC, this descriptive result does not independently establish that the cascade is unrelated to the report response.

### 2.3 Network Dynamics During Perceptual Switching

Having identified a distributed set of regions differentially engaged by spontaneous switching, we next asked how these regions relate to canonical large-scale brain networks—the review-derived BRAS network and the resting-state networks (RSNs)—and how those networks dynamically interact around the perceptual report.

#### Convergence with the review-derived BRAS network

We first compared our significantly modulated regions to the set of brain areas synthesised across studies in the review by Brascamp et al. [37]. We re-plotted these regions on a cortical surface using the HCP 360-area parcellation (Fig.7; hereafter the BRAS network). The dPC1-5 analysis (Fig.4A) recovered substantial regional overlap with the BRAS network, including areas 6r, 44, 45, FOP4, FOP5, AVI, LO3, MT, MST, FST, PGi, LIPv, 7PC, AIP, PF, PFt, PFop, TPOJ2, TPOJ3, FEF, 46, V6A, and DVT. A more selective subset— LO3, MT, MST, FST, and AIP—was retained in the dFC1 noise-correlation analysis (Fig.4B), which used fewer surrogates and therefore a less stringent per-map threshold but differed by removing the trial-averaged component, reinforcing the central role of motion-selective and intraparietal regions across the two analyses.

#### Structural overlap of BRAS with canonical resting-state networks

Beyond region-by-region overlap, prior work has shown that the BRAS network overlaps substantially with the canonical RSNs—particularly DAN and VAN [37]. We therefore examined the structural overlap of BRAS with eight RSNs derived from [47] (Fig.S2). Treating each network as a binary vector over the 758 MD758 regions and computing pairwise correlations after normalisation (Fig.5B) showed that BRAS overlaps most with DAN and VAN, less with the frontoparietal network (FPN), and is *negatively* correlated with the DMN. BRAS was also right-lateralised, evidenced by a positive correlation with a left–right (LR) lateralisation vector. Among the RSNs, the largest cross-network overlaps were consistent with prior reports: DAN with VIS, VAN with FPN and LAN, and SMN with AUD.

#### A temporal cascade of network engagement around the perceptual report

To resolve how each network engages dynamically around the perceptual report, we computed network-level PSTHs by projecting brain activity onto the standardised vector derived from each binary RSN mask at every time point, analogous to the region-level PSTHs in Fig.1(A,B). Because standardisation makes each vector sum to zero, these scores are inside-versus-outside spatial contrasts relative to the whole-brain mean and are insensitive to a spatially global component; the activation/suppression shorthand used below therefore does not denote absolute network activation. In the spontaneous condition (Fig.8A), the BRAS contrast showed a clear positive peak approximately 4 s post-report, occurring *concurrently* with a strong negative DMN contrast. The replayed condition (Fig.8B) yielded substantially weaker and temporally delayed BRAS modulation, and only a weak DMN response.

#### Network engagement relative to random area sets

To assess whether each RSN was engaged more than a random set of areas of the same size, we computed correlation z-scores between the task-differential dPSTH and each RSN, comparing against a null distribution generated by randomly shuffling RSN vectors across brain areas. This null does not preserve spatial autocorrelation, and no threshold or correction across time points and networks was applied, so these z-scores are best read descriptively. Both dSig (Fig.8C) and dPC1-5 (Fig.8E) showed positive BRAS and DAN contrasts together with a strong negative DMN contrast around 4 s post-report, with the dPC1-5 result notably sharper and more temporally focused; the VIS contrast was clear in dPC1-5 but weak and transient in dSig. Importantly, a similar post-report profile emerged in dFC1 (Fig.8G)—even though noise-correlation removes shared time-locked signals—with positive BRAS, DAN, and VIS and negative DMN scores between 3 and 8 s post-report. In dFC1, however, deviations of similar size were already present in the few seconds before the report (BRAS, DAN, and VIS around *z* ≈ 1.5–2; DMN reaching *z* ≈ −2), so the dFC1 profile is not specific to the post-report period. This suggests that the post-report profile is not fully explained by sensory or motor events that happen to co-occur with the report, though it does not exclude a contribution from trial-to-trial motor variability not removed by trial-mean subtraction (see Discussion). VAN and FPN also showed a more modest (*z* ≈ 1–2) and more sustained differential component that extended to around 10–14 s post-report, most evident in dSig (Fig.8C), consistent with the longer time scales captured by PC2 (Fig.3C); this later component is analysed in less depth here than the 4 s DAN–VIS–DMN complex and should accordingly be treated as a more preliminary part of the overall picture.

#### Functional decoupling of the DMN from task-positive networks

We also investigated the correlations between the dynamical evolutions of pairs of RSN networks. The RSN-by-RSN crosscorrelation matrices (Fig.8D,F,H) revealed a clear functional architecture beyond what structural overlap predicts. A striking and consistent feature of all three matrices was the negative correlation of the DMN with multiple task-positive networks—BRAS, DAN, VAN, SMN, and VIS— amplified in the denoised dPC1-5 matrix (Fig.8F). Positive correlations such as DAN–VIS were also more distinct in dPC1-5 than in dSig, while VAN–FPN was strong in both. In the dFC1 crosscorrelation matrix (Fig.8H), where most correlations were near zero, the strongest positive relationships were DAN–VIS, VAN–FPN, and BRAS–DAN/VIS, while the strongest negative relationships were DMN–DAN, DMN–VIS, and BRAS–DMN/LAN.

#### Functional and structural network architectures partially diverge

Comparing the task-based cross-correlations (Fig.8D,F,H) with the purely structural overlap matrix (Fig.5B) shows that some relationships—e.g., VAN–FPN and VIS–DAN—are consistent across both, while at least one diverges: BRAS and VAN overlap structurally (Fig.5B) but are barely correlated in the dFC1 matrix (Fig.8H), although they co-modulate moderately in dSig and dPC1-5. Based on this comparison, the functional architecture of perceptual switching appears not to be fully predicted by the anatomical layout of the resting-state networks, though we have not systematically quantified the proportion of network pairs for which structural and functional coupling agree versus diverge.

## 3 Discussion

This study mapped the spatiotemporal dynamics of brain activity associated with spontaneous versus replayed perceptual switching of a kinetic depth effect (KDE) stimulus, using a fine-grained MD758 parcellation, principal component analysis, and time-resolved noise-correlation functional connectivity. Three complementary analyses—the raw differential signal (dSig), the eigenvalue-weighted combination of principal components (dPC1-5), and the first noise-correlation component (dFC1)—revealed a distributed, temporally structured network response that distinguishes endogenous from externally driven perceptual transitions; as noted in Results, these are related but not fully independent views of the same underlying signal (dPC1-5 is a denoised transform of dSig, and dFC1 is computed from covariation among the same areas as dPC1), so their agreement should be read as convergent rather than as independent replication. From dSig we observed spontaneous-greater modulations 3–4 s post-report in parietal cortex, FEF, SCEF, the inferior frontal gyrus, and the frontal operculum; dPC1-5 sharpened and tightened this response, peaking at ~4 s post-report and, in the cortical-surface maps, also showing occipital areas; and dFC1, which by construction removes the trial-averaged motor component, identified bilateral MT and surrounding regions as central to the perceptual cascade. Inter-subject consistency was highest for dPC1-5 and clear for dSig, but only weakly elevated for dFC1; because ISC is computed on the same per-subject z-scored differential maps analysed elsewhere in this study, it should be read as evidence of consistent spatial pattern across subjects rather than as a fully independent replication that the cascade is not an artefact of the report-related button press.

### Distributed regional architecture: IFC, motion-selective occipital, parietal, and premotor cortex

The differentially activated regions span a coherent functional network. In the inferior frontal cortex (IFC), we found robust positive activation in IFG areas 44 and 45, frontal opercular areas FOP4 and FOP5, and anterior insula (AVI). These regions are consistently implicated in report-based studies of bistable perception [37, 48, 49], and prior reports that IFC activates earlier during spontaneous than replayed transitions [16] have been taken to suggest a role in inducing perceptual reorganisation [2], which our report-locked data cannot test directly. No-report work, however, indicates that part of this frontal activity may reflect introspection and reporting rather than the perceptual alternation itself [50], a caveat that applies equally to the present report-based design.

Occipital activation in LO3, MT, MST, and FST (Fig.4A,B) is consistent with the structure-from-motion nature of KDE: MT discriminates coherent from incoherent dot motion [51–55], integrates motion with binocular disparity [56, 57], and supports the recovery of three-dimensional structure from motion [29], while LO3 lies in lateral occipital cortex, a region situated between the dorsal and ventral streams that processes both shape and motion cues [58, 59]. In the parietal cortex, we recovered IPS (AIP, LIPv), SPL (7PC), IPL (PF, PFt, PFop, PGi), and TPJ regions (TPOJ2, TPOJ3), consistent with prior reports of parietal involvement in perceptual alternation [17, 26, 44, 61, 62] and in SFM-specific paradigms [27, 63–67]. Premotor activations (6r, FEF) and DLPFC area 46 further extend the network into circuitry supporting saccadic control [68–72], attention mapping, and goal-directed behaviour [37, 73–75]. SCEF, visible in the raw dSig but suppressed by PCA denoising, is an oculomotor and cognitive-control node whose presence in dSig but not dPC1-5 may reflect entanglement with the motor report.

### Principal components and resting-state network composition of the differential network

The two leading principal components reveal an additional temporal structure that complements this regional inventory: PC1 captures a stereotyped global cycle of pre-report suppression, a sharp post-report peak around 4 s, and a below-baseline rebound near 10 s (Fig.3A), while PC2 carries more sustained, hemispherically asymmetric modulations between 4 and 10 s post-report (Fig.3C). The first component may partly reflect the attentional cascade we describe below, although it may also carry report-related motor variance (see Results), whereas the second may index slower lateralised cognitive-control processes that unfold once the perceptual transition is established. At the level of canonical resting-state networks, the regions we recovered map onto specific HCP-defined nodes of the DAN (7PC, AIP, FEF, LIPv, MST, MT [95, 97]) and VAN (6r, AVI, FOP4, p9-46v, PGi, TPOJ2 [96]), with TPOJ2 and PGi showing a left-lateralised pattern that contrasts with the predominantly right-lateralised VAN activation typically reported in the bistable-perception literature.

### A pre-report reduction in activity during spontaneous switching

A further feature of our data not captured by the post-report peak alone is that activity averaged across brain areas was reduced in the few seconds *before* the perceptual report in the spontaneous, but not the replayed, condition (Fig.1C), although no individual area showed a pre-report difference beyond what is expected by chance (Fig.2A). The trial-by-trial standard deviation of activity was also more strongly and more rapidly suppressed around the post-report peak in spontaneous switching (Fig.1D,F), indicating that the post-switch dynamical trajectory was highly stereotyped across trials. Together, these two observations are consistent with the view that a spontaneous perceptual switch is preceded by a transient endogenous reorganisation of cortical activity—an “anticipatory preparatory state”—that then resolves into a reproducible post-switch trajectory. One possible interpretation is a slow accumulation process before attractor crossing in mutual-inhibition models of bistable perception [6–8]; the present data do not establish this mechanism. The pre-report suppression was evident only in the spontaneous condition, and the post-report variability suppression, although also present in the replayed condition, was weaker and later there (Fig.1F); this pattern is consistent with endogenous, internally generated reorganisation, but a condition-by-time interaction was not formally tested. Moreover, spontaneous transitions were self-timed and temporally unpredictable, whereas replayed transitions were externally timed and predictable in their statistical structure, and the two conditions used different ring geometries (see Methods). The dissociation could therefore reflect temporal-expectation, motor-timing, or low-level stimulus confounds, or differences between conditions in the overlap of responses to neighbouring reports (see Limitations), rather than a specifically perceptual anticipatory process; disentangling these possibilities would require controls that vary temporal predictability and stimulus geometry independently of perceptual origin.

### A cascade of large-scale network dynamics follows the resolution of perceptual ambiguity

Beyond the regional architecture, our network-level analyses revealed that spontaneous perceptual switching is followed by a dynamic cascade of engagement across resting-state networks. We use “cascade” here in a descriptive sense, to denote a temporally ordered sequence of network engagement; our design, based on report-locked correlational measures, does not establish causal or mechanistic dependence between its stages, and the two stages described below were comparatively weakly coupled in the dFC1 analysis, without a formal test of that specific coupling (see below). Around 4 s post-report we observed concurrent positive BRAS, DAN, and VIS contrasts and a negative default mode network (DMN) contrast, with VAN and FPN engaging later, and more modestly, around 10 s (Fig.8). The early BRAS–DAN–VIS–DMN profile also emerged in dFC1, despite its removal of the trial-averaged time-locked component (although there it was not confined to the post-report period), indicating that this profile is not fully explained by time-locked sensory or motor events, though—as with the regional analyses above—this does not rule out a contribution from residual motor variability not captured by the trial mean. This is consistent with the broader picture that endogenous brain activity dynamically transitions between large-scale network states [98–100], with such transitions known to underlie working memory, cognitive control, and attentional engagement. The DMN– task-positive anti-correlation we observed (Fig.8F,H) is consistent with the broader literature on DMN suppression during attentionally demanding tasks [77, 78], and with clinical work linking the DMN to consciousness and awareness [79, 80]. The behavioural relevance of such anti-correlation is supported by attention studies: stronger pre-trial DMN–task-positive anti-correlation predicts faster behavioural responses in psychomotor vigilance [90], and pre-stimulus network connectivity patterns predict conscious perceptual detection [91], raising the possibility that the stronger negative DMN contrast we observe in the spontaneous condition is functionally related to the concurrent positive BRAS contrast; testing a facilitatory relationship directly within the present dataset (e.g., via trial-level covariation between DMN contrast depth and BRAS response magnitude) remains for future work.

### Functional network architecture diverges from anatomical overlap

Comparison of the functional cross-correlation matrices with the structural overlap matrix (Fig.5B) suggests that the functional architecture of perceptual switching is only partially predicted by anatomical network layout. Some couplings track structural overlap closely—most notably VAN–FPN and DAN–VIS, which are both consistent across the two views—while at least one diverges: BRAS and VAN overlap structurally but are barely correlated in the dFC1 matrix during perceptual switching. This suggests the task-related functional architecture is not simply a passive reflection of the resting-state network layout, though we have not systematically characterised what proportion of network pairs follow this divergent pattern versus the structurally-consistent one.

### Convergence with review-derived findings and novel temporal dissection

The differentially activated regions show substantial regional overlap with the review-derived BRAS network synthesised by Brascamp et al. [37] (this was solely a qualitative, region-by-region comparison against the plotted BRAS coordinates; we did not compute a formal overlap statistic such as a Dice or Jaccard coefficient against a null distribution, which we identify as a useful addition for future work). Overlap included all of the canonical right-lateralised frontoparietal regions (IFC, FEF, DLPFC, IPS), as well as MT. The BRAS mask itself was structurally right-lateralised, as indicated by its positive correlation with the left–right vector in Fig.5B, and the dominant inferior-frontal/anterior-insular components in our differential maps emerged predominantly in the right hemisphere (Fig.4A), consistent with the classical right-hemispheric bias reported across multiple bistable-perception studies [16, 25, 37, 44]. What our analysis adds is a report-locked temporal dissection of this network at second-scale resolution. By projecting the differential signal onto resting-state networks, our results suggest that the BRAS response is not a unitary activation but rather a composite of (i) an early, sharp DAN–VIS coactivation coupled with DMN suppression around 4 s post-report, and (ii) a more modest, sustained VAN–FPN differential component extending to around 10–14 s post-report, most evident in dSig and treated here as exploratory; these two components were comparatively weakly coupled in the dFC1 cross-correlation matrix (Fig.8G,H), where BRAS–FPN and BRAS–VAN did not appear among the strongest relationships, though we did not formally test these specific couplings against a null distribution. This decomposition is broadly consistent with, and adds temporal precision to, recent network-level studies of bistable perception. Mao et al. [76] related the temporal variability of resting-state (task-free) functional connectivity to individual differences in average percept duration during a separate bistable task, reporting that perceptual transitions depend on coordinated DMN, FPN, cingulo-opercular (CON), DAN, and salience network (overlapping with VAN) activity, with DAN–SN and DAN–CON variability predicting percept duration; this is a trait-level association rather than a report-locked, within-switch temporal decomposition. Karten et al. [81] used block-based psychophysiological interaction analysis to describe dynamic LOC–DMN–FPN coupling that distinguished sustained dominant versus alternative percept states, rather than resolving the sub-10-second temporal structure around an individual switch. Relative to both, the present report-locked PSTH approach resolves network engagement with second-scale precision around the moment of the perceptual switch itself. Our results agree with the central role of cooperative DAN activity in these studies, and additionally highlight a tight DAN–VIS coupling and an early DMN suppression that accompany the network-level reorganisation following a spontaneous perceptual transition. The tight DAN–VIS coupling we observe is consistent with established causal influences from the DAN onto visual cortex during attentionally demanding tasks [43, 92–94]; in particular, Spadone et al. [92] showed that visuospatial attention enhances DAN-driven top-down input to specific intermediate visual areas (V3a–V7 and MT) via FEF and IPS/SPL—the same nodes we recover here as central to the dPC1-5 cascade. The DAN–VIS coupling may therefore reflect attentional up-regulation accompanying or following a spontaneous perceptual transition.

### Methodological strengths and limitations

The combination of high-resolution cortical parcellation (MD758), PCA-based denoising, and time-resolved noise-correlation functional connectivity proved useful for reducing the contribution of the strong stereotyped motor response associated with the button-press report, though several limitations qualify how far this reduction, and the cascade findings more broadly, can be interpreted. First, dFC1, dPC1-5, and dSig are related transforms of the same underlying signal rather than fully independent analyses (see Results and Discussion above); their agreement is best read as internal convergence rather than independent replication, and the noise-correlation approach removes only the trial-averaged motor component, not trial-to-trial motor variability or condition-level differences in motor-timing predictability. Second, because all analyses are locked to the perceptual report rather than to the switch itself, and given typical haemo-dynamic response lags and unreported reaction-time variability between the covert switch and the overt report, the temporal ordering we describe (an early DAN–VIS–DMN complex followed by a later, more modest VAN–FPN component) is best read as a description of report-locked activity rather than a validated causal or mechanistic sequence; for spontaneous switches the covert switch time is unobservable, and although reaction times to the physical reversals could be derived for the replayed condition, they were not analysed here. Third, our cohort comprised only six participants, which limits the generalisability of both the precise quantitative parameters and the qualitative cascade description reported here; some analyses (e.g., the cortical maps in Figs.2 and 4) pool across the six subjects into a single combined-subject test rather than treating subjects as a random effect, which is a stronger statistical claim about this specific sample than about the population from which it was drawn. Fourth, the spontaneous and replayed conditions used symmetric and asymmetric ring geometries, respectively, so every condition contrast confounds perceptual-switch origin with low-level stimulus geometry and cannot uniquely attribute a difference to endogenous switching. Fifth, the standardised RSN projections are inside-versus-outside contrasts that remove spatially global components; positive and negative scores therefore cannot be interpreted as absolute network activation and deactivation. Sixth, the mapping from resting-state network identity (DAN, VAN, DMN) to the cognitive processes conventionally associated with them (e.g., attentional control, internally directed thought) is inferred from prior literature rather than directly tested in the present task. Seventh, although eye position was recorded, eye movements were not analysed; because MT, MST, FEF, and SCEF are also central to smooth pursuit and saccade control, and rotating displays can drive pursuit and optokinetic eye movements, differences in eye movements between conditions or around switches could contribute to the differential maps. Eighth, the report-locked PSTHs were obtained by averaging rather than by deconvolution; because inter-report intervals were often shorter than the ± 15-s analysis window (Fig.S3), each window can contain responses to neighbouring reports, and differences between conditions in the timing of those reports could contribute to small differential effects such as the pre-report reduction. Future work with larger samples, better-matched control stimuli, analysis of eye movements, deconvolution-based response estimates, explicit reaction-time modelling, and complementary methods such as dynamic causal modelling and multimodal (EEG– fMRI) imaging will be important to establish the population-level robustness of the cascade and to test the directionality of the network dynamics we describe.

### Towards a phenomenological link

An open question raised by the present work, though not addressed by our data, is whether the distinct neural profiles of spontaneous versus replayed switching correspond to distinct subjective experiences of the two transition types. Phenomenological work using binocular rivalry has begun to dissociate the subjective signatures of spontaneous versus stimulus-driven perceptual transitions [87]; extending a comparable approach to KDE—noting that rivalry and KDE are phenomenologically distinct paradigms, so this extension is not immediate— could in principle test whether the network profile we identify here covaries with specific phenomenological features of the spontaneous switch [88, 89].

In summary, by combining a fine-grained parcellation with principal-component and noise-correlation analyses, we have shown that spontaneous perceptual switching of a bistable KDE stimulus is followed by a dynamic cascade of large-scale network dynamics: an early, sharp positive dorsal-attention and visual-network contrast together with a negative default-mode-network contrast, followed by a delayed and more modest VAN–FPN component. The early profile appears in all three complementary analytic measures and is only partially dissociable from the report-related motor response, subject to the limitations discussed above. Beyond converging with and refining the review-derived BRAS network, our results foreground network dynamics, rather than isolated regional activations, as an important contributor to the network-level reorganisation that follows resolution of perceptual ambiguity in this paradigm.

## 4 Methods

### 4.1 Experimental methods

#### 4.1.1 Participants

Six participants (two female and four male, 24–35 years old) participated in the experiment. All participants were healthy individuals with normal or corrected to normal vision. During this experiment, we followed the ethical guidance of the Centre for Neuroscientific Innovation and Technology, Magdeburg. Participants in functional imaging studies were medically screened and gave written informed consent to neuroimaging according to the guidelines of the Centre for Neuroscientific Innovation and Technology (ZENIT), Magdeburg. Ethics approval for the proposed study of multistable perception, including recording of eye movements, was obtained from the Ethics Commission of the Faculty of Medicine of the Otto-von-Guericke University, Magdeburg, under permit number “29/21 No-report monitoring of subjective visual perception” and “14/23 Kortikale Auflösung des spontanen Kippens der visuellen Wahrnehmung.” Participants gave separate written informed consent for the behavioural aspects of the study. Consent forms were approved by the Data Protection Office of the Faculty of Informatics, Otto-von-Guericke University, Magdeburg.

Participants sat or lay comfortably and viewed visual displays through 45-degree mirrors. Eye position was recorded by an infrared camera through the mirrors (which are transparent to infrared light). Experimental runs lasted up to 6 min, training sessions up to 90 min per day (including breaks), and imaging sessions up to 150 min per day (including breaks and time for setting up eye tracking). Participants were instructed to attentively watch the display and to report changes by a manual button press. They were free to rest between runs and were invited to participate in no more than 12 sessions distributed over four weeks. They were free to abort runs and sessions at any time, without financial penalty. They were free to leave the study at any time, again without financial penalty. The standard rate for participant payment was C12 per hour. After participating in 12 sessions of 90 min duration, participants could earn up to C216 in total.

#### 4.1.2 Stimuli

Stimuli were generated using Psychtoolbox-3 [82] running in MATLAB (Mathworks, Natick, MA, USA), and presented in two conditions designed to elicit spontaneous and replayed reversals, respectively. The first stimulus consisted of two coaxial dotted rings forming a 60° visual angle rotating in the frontal plane around a vertical axis (Fig.9). To prevent observers from focusing on local visual patterns instead of observing the whole object, only a certain portion of the dots in each ring were shown, and they were slowly replaced by other dots during the presentation.

The second stimulus was made up of two asymmetric bent rings with visual characteristics similar to the first stimulus (e.g. number of dots, width, and thickness). To reduce the frequency of spontaneous perceptual reversals, we introduced asymmetry to the rings. From this point, we refer to the stimuli accompanying spontaneous and replayed reversals, as symmetric rings and asymmetric rings, respectively.

#### 4.1.3 Procedure

First, participants attended two training sessions, which consisted of four runs of the symmetric rings followed by two runs of the asymmetric rings. In order to generate replayed reversals temporally similar to the spontaneous reversals, enough data on temporal distribution of the spontaneous reversals was needed. Therefore, the number of runs with the symmetric rings was higher than the number of runs with the asymmetric rings during training sessions. Later, each participant attended two fMRI scanning sessions, with two runs each of the symmetric and asymmetric rings per session. Each run lasted 330 s.

To reduce possible perceptual biases resulting from previous visual memory, each block contained 30 s of initial priming, during which the direction of rotation of the rings changed every 5 s. In addition, the luminance of the objects was manipulated to ensure the perception of depth, by dimming the backside of the stimuli. Observers were guided to react whenever they observed the rotating rings change their direction of rotation. Therefore, the priming phase also served to confirm that observers understood the instructions. After priming, the symmetric rings kept rotating without changing their direction of rotation. Therefore, any reported reversals were spontaneous reversals. In contrast, asymmetric rings were forced to change their direction of rotation by the temporal distribution of Γ(*μ, CV*) of the coefficient of variation *CV* = 0.8 [83] and the average of reversal time measured during the symmetric rings presentations.

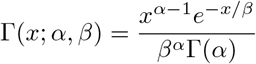

where Γ(*α*) is the gamma function and *α* = 1*/*(*CV*)^2^, *β* = *μ*(*CV*)^2^.

#### 4.1.4 fMRI acquisition

A 3T Siemens Prisma scanner with 64-channel head coil was used for image acquisition. Structural T1-weighted images were acquired with a standard MPRAGE sequence (1 mm isotropic voxels, TR = 2500 ms, TI = 1100 ms and TE = 2.82 ms). T2*-weighted sequence (3 × 3 × 3.6 mm, TR = 1000 ms, TE = 30 ms) was used for functional images. Field maps were acquired by dual echo gradient sequences (1.594 × 1.594 × 2 mm, TR = 720 ms, TE1 = 4.92 ms, TE2 = 7.38 ms).

#### 4.1.5 fMRI preprocessing

Preprocessing steps were the same as those in [84]. Briefly, DICOM files were converted into nifti (MRIcron, NIH, Bethesda, MD, USA), and brain extraction, anatomical image segmentation, fieldmap correction, motion correction, spatial smoothing, high-pass temporal filtering, and registration to structural images and the standard MNI152 2 mm template were done in FSL software (FMRIB, Oxford, UK). Additionally, every voxel’s activity was despiked using BrainWavelet toolbox [85]. Lastly, we regressed out the mean CSF activity as well as 12 DOF translation and rotation factors predicted by motion correction algorithm (MCFLIRT).

### 4.2 Data analysis

Voxels were clustered into 758 brain areas according to the MD758 parcellation [84]. This yielded four fMRI runs of 300 s each for each of six subjects in two task conditions, spontaneous and replayed perceptual switching (each 330 s run consists of a 30 s priming period followed by the 300 s of stimulation analysed here; the four runs of a given condition were acquired as two runs in each of the two scanning sessions). We denote these time series *V* ^*T Sβ*^(*A, t*) where *A* = 1, ‥, 758 labels the brain area, *t* = 1, …, 300, the time elapsed in seconds, *T* the task condition, spontaneous or replayed, denoted by superscripts ‘spnt’ and ‘frcd’ respectively (‘frcd’ from ‘forced’, an earlier label for the replayed condition retained in the notation), *S* = 1, ‥, 6 the subject, and *β* = 1, ‥, 4 the run. First *V* ^*T Sβ*^(*A, t*) are standardised for each area *A* by subtracting the mean across time, *t*, and dividing by the standard deviation across time, *t* to obtain *X*^*T Sβ*^(*A, t*).

All results in this paper proceed by constructing report-locked peri-switch time histograms (PSTHs). Perceptual reports are referred to as ‘trials’ in the following. In the simplest case these PSTH are calculated from *X*^*T Sβ*^(*A, t*) itself. In other cases however *X*^*T Sβ*^(*A, t*) is first transformed to another quantity, *Q*^*T Sβ*^(*A, t*), before PSTH are constructed. In each run, *β*, for each subject *S* in each task, *T*, the *n*^th^ perceptual switch report time is denoted 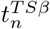.We do not include in 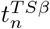 perceptual reports within 16 s of the start or end of a run. Perceptual report intervals are described in Fig.S3. To obtain cross-trial averages of any given quantity, *Q*^*T Sβ*^(*A, t*), 31 s segments centred on 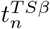 are selected, 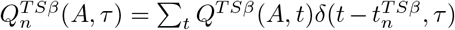,where *δ*(*x, y*) is the Kronecker delta and *τ* = −15, ‥, +15. These are then averaged, 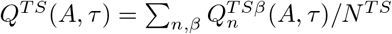 where 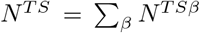 is the number of report trials across all runs *β* for subject *S* in task *T*. The operation of cross-trial averaging going from *Q*^*T Sβ*^(*A, t*) to *Q*^*T S*^(*A, τ*) is denoted ⟨*Q*^*T Sβ*^(*A, t*)⟩_tri_.

We also obtain cross-trial standard deviations, 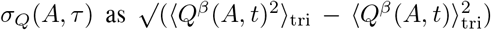 and cross-trial noise correlation matrices, *σ*_*Q*_(*A*_*i*_, *A*_*j*_, *τ*) = (⟨*Q*^*β*^(*A*_*i*_, *t*)*Q*^*β*^(*A*_*j*_, *t*)⟩_tri_ − *Q*(*A*_*i*_, *τ*)*Q*(*A*_*j*_, *τ*))*/*(*σ*_*Q*_(*A*_*i*_, *τ*)*σ*_*Q*_(*A*_*j*_, *τ*)).

For presentation simplicity trial averaged quantities are denoted *Q*(*τ*) in contrast to original time series which are denoted *Q*(*t*). Fig.1(A) shows *X*^spnt^(*A, τ*) (referred to as Sig(*A, τ*) in the text) obtained by averaging the brain activation PSTH, *X*^spnt,*S*^(*A, τ*), over subjects, *S*. Fig.1(B) shows *X*^frcd^(*A, τ*). Fig.1(D) shows 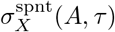 obtained by averaging the 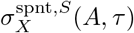 over subjects, *S*. Although all calculations are performed using the 758 brain areas of the MD758 brain parcellation [84], for visualisation purposes the results are averaged down to the 90 automated anatomical labeling (AAL) parcellation [86]. Fig.1(C,F) show these quantities further averaged over brain areas *A*.

We also calculate transformations to the original *X*^*T Sβ*^(*A, t*) before applying the cross-trial averaging procedure described above. To obtain the principal components, denoted ‘PC’, first runs, *β* = 1, ‥, 4, are combined to produce 300 × 4 observations of 758 areas *A* for each subject *S* in the spontaneous task, denoted *Y* ^*S*^(*A, t*), *t* = 1, ‥1200. These are again standardised by subtracting the mean across *t* and dividing by the standard deviation across *t* for each area *A*. The 758 × 758 cross-correlation matrix, *Y Y* ^*′*^ is calculated, (*Y* ^*′*^ denotes *Y* transpose) and its first five eigenvectors obtained, denoted 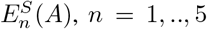 and associated eigenvalues 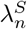.Using these eigenvectors new area activations 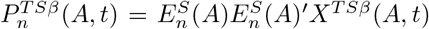 are obtained. Eigen value weighted sums, 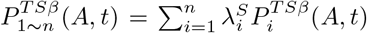 are also calculated. Finally cross trial averages 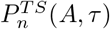 and 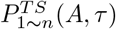denoted ‘PCn’ and ‘PC1-n’ in the text, respectively, are obtained as described above.

Fig.3(A,C) show 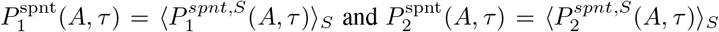 obtained by averaging over subjects *S* and further averaging into the 90 AAL brain areas in the spontaneous perceptual switching condition.

To obtain differential PSTH z-scores we subtract replayed task PSTH from spontaneous ones to obtain differential PSTH (dPSTH), *dQ*^*S*^(*A, τ*) = *Q*^spnt,*S*^(*A, τ*) − *Q*^frcd,*S*^(*A, τ*) and also combined subject versions, *dQ*(*A, τ*) = ⟨*dQ*^*S*^(*A, τ*)⟩_*S*_. We then calculate z-scores, by comparing the given value *dQ* with surrogates *dQ*_*surr*_. A surrogate is created by resampling perceptual report trials from the pooled spontaneous and replayed task conditions. *N* ^spnt,S^ spontaneous task trial segments and *N* ^frcd,S^ replayed task trial segments are drawn randomly with replacement from the full set,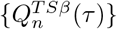,including all runs *β*, both tasks *T* and all trials *n* for a given subject *S* to calculate a pair of surrogates 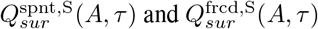 and an observation of 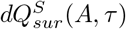 and *dQ*_*sur*_(*A, τ*).

From the surrogates we calculate a symmetrical ‘p-score’, *dQ*_*p*_ = ((*α*(*N*_sur_−1)*/N*_sur_)+1)*/*(*N*_sur_+ 1), where *α* =∑ _*sur*_ Θ(*dQ* − *dQ*_*sur*_) and Θ(…) is the Heaviside function, and *N*_sur_ is the number of surrogates. When the true observation is larger than all the surrogates this quantity is close to unity, and when the true value is smaller than all the surrogates it is close to zero. (Note that this p-score calculation is similar to that used to obtain a one-sided p-value (*p* = (*N*_sur_ −*α*+1)*/*(*N*_sur_ +1)) but not identical because it is symmetrical and 0 *< dQ*_*p*_ *<* 1.) We further log transform it to calculate logarithmic z-scores (denoted dPSTH_*z*_), according to the ‘surprise’, *dQ*_*z*_ = *Z*(*dQ*) = log(*dQ*_*p*_) − log(1 − *dQ*_*p*_) which is large and positive when the true observation is larger than all the surrogates and large and negative when the converse is true. Note this definition of the z-score is also different to the standard one, (which is given by *dQ*_*z*_ = (*dQ* − ⟨*dQ*_surr_⟩)*/σ*(*dQ*_surr_) depending on the mean and standard deviation of the *dQ*_surr_ surrogates 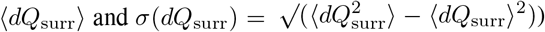 However it is less susceptible to large deviations which may occur. In fact results calculated using the standard definition are almost identical.

#### Multiple comparisons

Aside from these p-score and z-score calculations we also use the surrogates directly to provide the true p-values, *p* = 1*/*(*N*_sur_ + 1) for *α* = *N*_sur_ and *p* = 2*/*(*N*_sur_ + 1) for *α* = *N*_sur_ − 1, shown in Figs.2,4 and Supplementary Fig.S1, for non-parametric, distribution-free significance testing for each (brain area, time point) pair. Because we evaluate 758 brain areas at each of 31 time points relative to the perceptual report (758 × 31 = 23,498 tests), the family-wise error rate is non-trivial: at the strictest per-test level shown, *p* = 1*/*(*N*_sur_ + 1) (see below), 23,498*/*3000 ≈ 7.8 false positives are expected per map, when *N*_sur_ = 3000, by chance alone considering one tail (approximately double this, ~16, when both the positive and negative tails are counted together, and roughly double again, ~31, when the yellow (*α* = *N*_sur_ −1) and cyan (*α* = 1) levels in Fig.2(A,B) are also counted), and correspondingly more (~47–94) for the noise-correlation analyses evaluated against *N*_sur_ = 500 surrogates. Throughout the figures we therefore visualise *extreme* significance levels—specifically the subset of regions for which the observed differential PSTH exceeded *all* (or, for dSig and dPC1-5, all but one) of the *N*_sur_ = 3000 surrogates (all of *N*_sur_ = 500 for the more computationally demanding noise-correlation analyses)—which corresponds to per-test levels of *p* = 1*/*(*N*_sur_ +1) or 2*/*(*N*_sur_ +1) and is substantially more stringent than conventional *p <* 0.05, though as the calculation above shows, not sufficient on its own to reduce the expected false-positive count to zero. Convergence across the three complementary analyses (dSig, dPC1-5, and dFC1), together with the inter-subject consistency analysis (Fig.6), provides some additional protection against false positives arising from any single comparison; however, these three analyses are related transforms of the same underlying signal rather than fully independent tests (see Discussion), so this should not be considered equivalent to formal family-wise-error or falsediscovery-rate correction.

Fig.1(E) shows 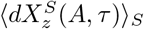 and Fig.3(E) shows 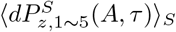 which are calculated from subjects separately and then averaged over subjects. Fig.2 shows dPSTH p-values calculated from the combined subjects for (A) the raw signal, *dX*_*p*_(*A, τ*) and (B) the PC1-5 method, *dP*_*p*,1*~*5_(*A, τ*), for brain areas with *α* = *N*_sur_ (red), *α* = *N*_sur_ − 1 = 2999 (yellow), *α* = 0 (blue) and *α* = 1 (cyan) where *N*_sur_ = 3000. Fig.S1 shows *dX*_*p*_(*A, τ* = 3) and Fig.4A shows *dP*_*p*,1*~*5_(*A, τ* = 4) for brain areas with *α* = *N*_sur_ (red) and brain areas with *α* = *N*_sur_ − 1 = 2999 (yellow) where *N*_sur_ = 3000.

Noise correlation functional connectivity measures are calculated from standardised time series *X*^*T Sβ*^(*A, t*) as follows. First noise correlation matrices, 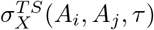 are obtained as described above. The first five eigenvectors and eigenvalues are obtained from this matrix at each time 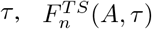, denoted ‘FCn’ in the text, and 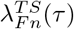. Fig.3(B,D) show 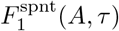 and 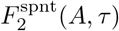 respectively, obtained by averaging 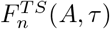 over subjects *S* and the 90 AAL areas in the spontaneous task condition.

In fact it is an issue with principal component analysis that the sense of the component is not defined; if 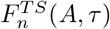, is an eigenvector so is –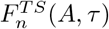. We assigned the sense of each eigen-vector at each time *τ* so that dot products with the eigenvectors at other *τ* were positive as far as possible. We are free to multiply the eigenvectors at different *τ* by an arbitrary factor of ±1.

We choose this factor as follows. First we find the correlations between eigenvectors at different τ,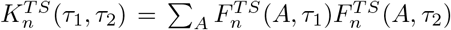 Next we calculate 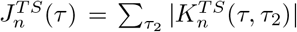 where |.| denotes the absolute value. Then we find 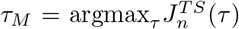, the eigenvector which has the largest absolute correlation with the other eigenvectors. Next we multiply eigenvectors 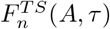at each *τ* by the factor 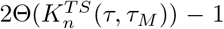, where Θ(.) is the Heaviside function.

This operation is repeated for each task *T* and subject *S*. Subsequently the operation is performed across subjects, where we now calculate 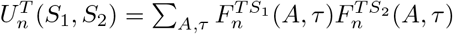, find 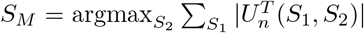 and multiply 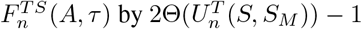. Finally the operation is performed across tasks *T*, to obtain 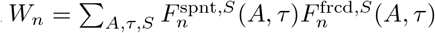 and the replayed-task eigenvectors 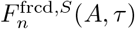 are multiplied by 2Θ(*W*_*n*_) − 1, with the spontaneous task as reference.

In principle this procedure could generate spurious correlations even between random vectors. However this is not a problem because all results reported are z-scores constructed by comparison with surrogates (except for time series visualisation figures) and this alignment procedure was always also applied to surrogates.

We again calculate task-differential dPSTH from the spontaneous and replayed 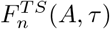, as 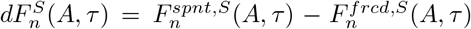 for each subject and also surrogates of this quantity to obtain z-scores as described above. When eigenvectors are obtained from surrogates the same alignment operations are always applied to the surrogates. In addition we align each given differential surrogate 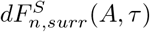 with the original dPSTH 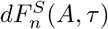 before calculation of z-scores.

Fig.3(F) shows dPSTH z-scores from the first FC component, 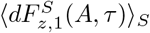, for subjects separately before averaging. Fig.2(C) shows dPSTH p-values calculated from the combined subjects for the first FC component *dF*_*p*,1_(*A, τ*) for brain areas with *α* = *N*_sur_ (red) and *α* = 0 (blue) where *N*_sur_ = 500. Fig.4B shows *dF*_*p*,1_(*A, τ* = 4) for brain areas with *α* = *N*_sur_ (red) and brain areas with *α* = 0 (blue) where *N*_sur_ = 500.

In Figs.S1, 4A, 4B and 7, we label brain areas using the 360 area parcellation of Human Connectome Project (HCP) [60], with region labels in the figures following the connectomic atlas nomenclature of [38], taking advantage of the parcellation for use in Freesurfer (https://surfer.nmr.mgh.harvard.edu/fswiki).

Fig.6 shows the mean inter-subject consistency (ISC). To calculate ISC values, differential PSTH z-scores 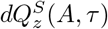 for each subject are centred and scaled across areas *A* at each time *τ* for each subject *S*, by subtracting the mean across *A* and dividing by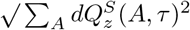 to obtain 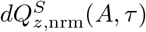. At each time *τ*, pairwise spatial similarity scores 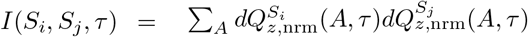are obtained. ISC(*τ*) shown in Fig.6 is 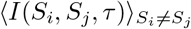, the mean across the 15 unique subject pairs. Because the scaling norm is computed before centring, *I* is the descriptive score defined here rather than Pearson’s *r* or standard cosine similarity.

**Figure 6.**
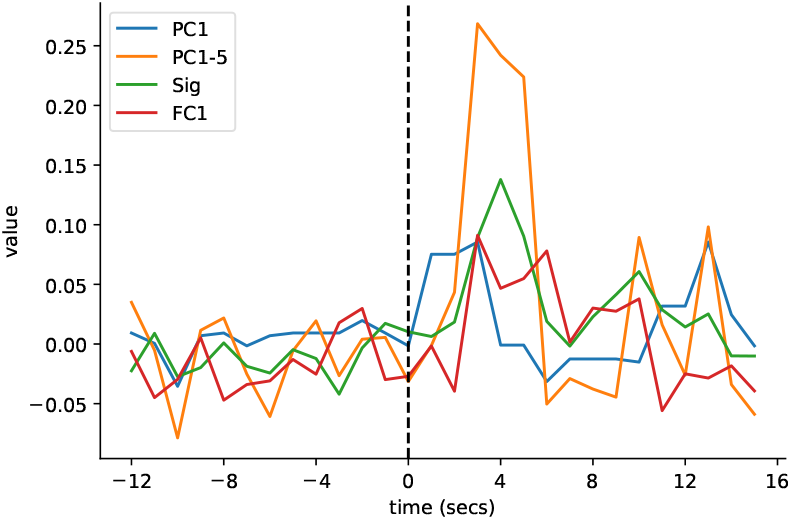
Mean inter-subject consistency (ISC) calculated from individual subject pairs. Line shows mean across subject pairs. Results shown for the first principal component (dPC1, blue; labelled “PC1” in the figure legend), the eigenvalue-weighted sum (dPC1-5, orange; “PC1-5”), the raw signal (dSig, green; “Sig”), and the first noise-correlation component (dFC1, red; “FC1”). The “d” prefix used in the main text denotes the task-differential measure computed from each base quantity (see Methods).

To obtain activations in Fig.8(A,B) we calculate correlations of the raw signal *X*^*T Sβ*^(*A, t*) with resting-state networks (RSN) [47], depicted in Fig.S2, the Brascamp areas (BRAS) [37] shown in Fig.7, and the lateralisation (LR) network. Any particular network is represented as length 758 binary vector of ones and zeros, denoted *R*(*A*). The LR network has ones on the right hemisphere. *R*(*A*) is standardised across areas *A* by subtracting its mean and dividing by its standard deviation across *A* to obtain *R*_nrm_(*A*). The scalar product *X*^*T Sβ*^(*R, t*) = *X*^*T Sβ*^(*A, t*)*R*_nrm_(*A*)^*′*^ is obtained. Since *R*_nrm_(*A*) sums to zero, this product is an inside-versus-outside network contrast relative to the whole-brain mean and is insensitive to a spatially global component. Here *X*(*R, t*) means the signal over networks *R* instead of areas *A*. Then we apply the cross-trial averaging to obtain *X*^*T S*^(*R, τ*) = ⟨*X*^*T Sβ*^(*R, t*)⟩_tri_. Fig.8(A,B) shows the average and SEM of *X*^*T S*^(*R, τ*) across subjects separately for spontaneous and replayed perceptual switching.

**Figure 7.**
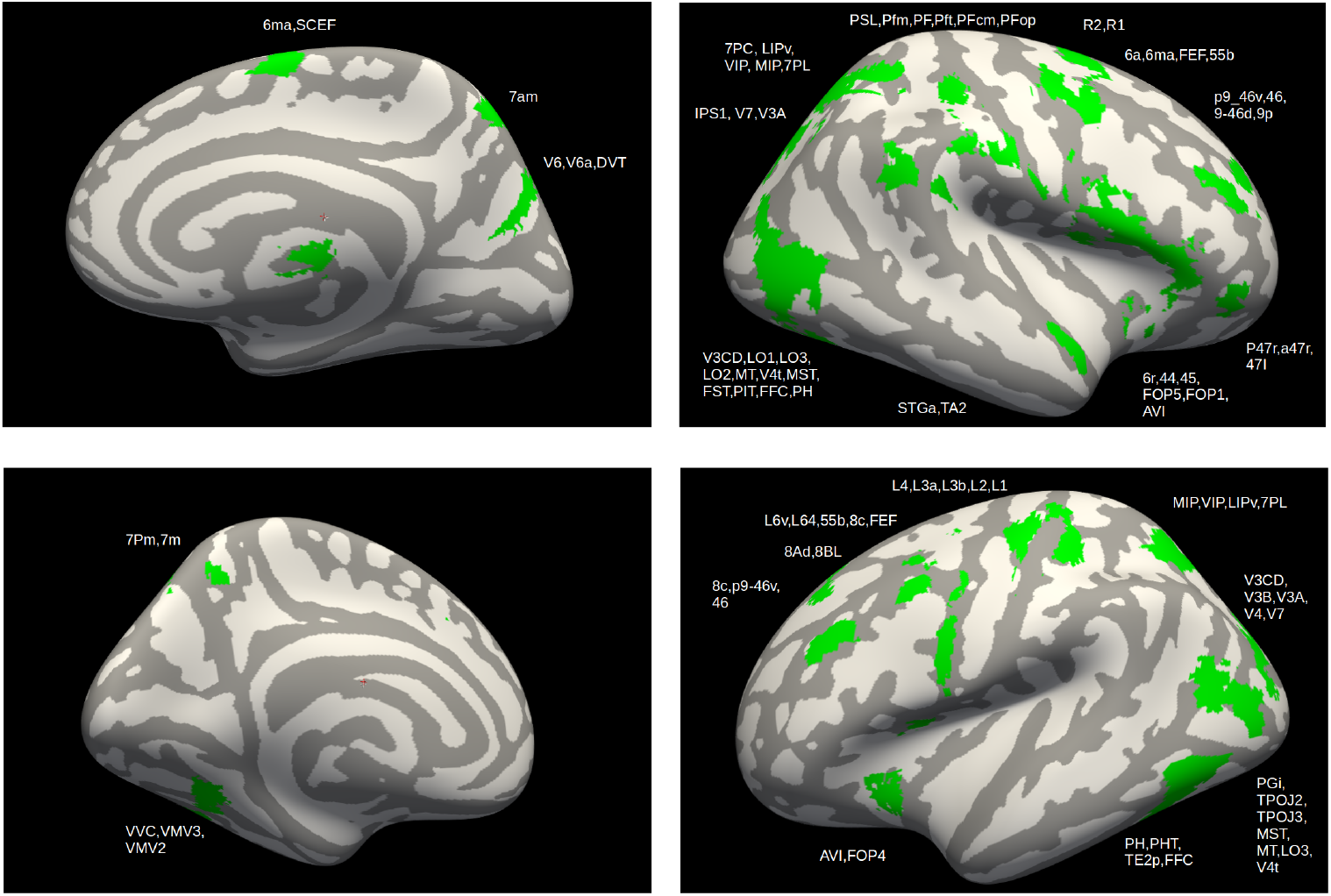
BRAS network areas implicated in spontaneous perceptual reversal across multiple studies as synthesised in the review by [37]. The review describes these areas as showing higher activation in spontaneous compared to replayed perceptual switching. We obtained the published coordinates and re-plotted them on a brain surface, labelled according to the HCP 360-area parcellation [38, 60].

**Figure 8.**
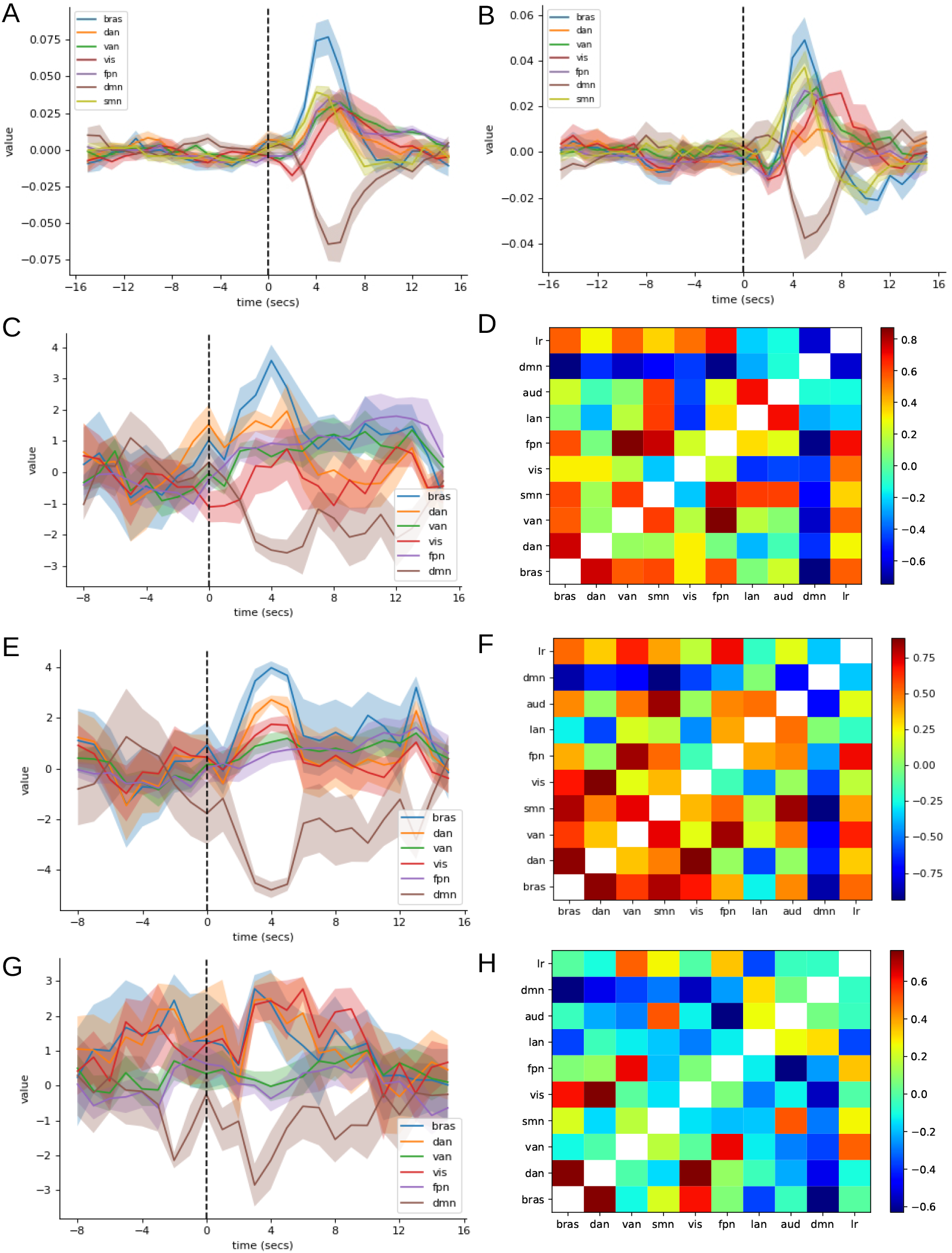
PSTH locked to perceptual report at *τ* = 0 of BRAS and RSN network contrast scores in (A) spontaneous and (B) replayed perceptual switching, calculated for individual subjects. Solid lines: mean across subjects; shading: SEM across subjects. (C,E,G) z-scores of correlation between task-differential dPSTH and RSNs for (C) the raw signal dSig, (E) dPC1-5, (G) dFC1. Per-subject z-scores then averaged; bands show SEM. (D,F,H) Cross-correlation matrices between task-differential dPSTH for different RSNs, computed from (D) dSig, (F) dPC1-5, (H) dFC1, corresponding respectively to (C), (E), (G). Combined-subject results. Network labels in the legends and matrix axes are shown in lowercase (*bras, dan, van, vis, fpn, dmn, smn, lan, aud, lr*) for visual compactness; these correspond to the BRAS, DAN, VAN, VIS, FPN, DMN, SMN, LAN, AUD, and LR networks as named in the main text. The SMN trace is included in (A,B) but omitted from (C,E,G) for visual clarity.

**Figure 9.**
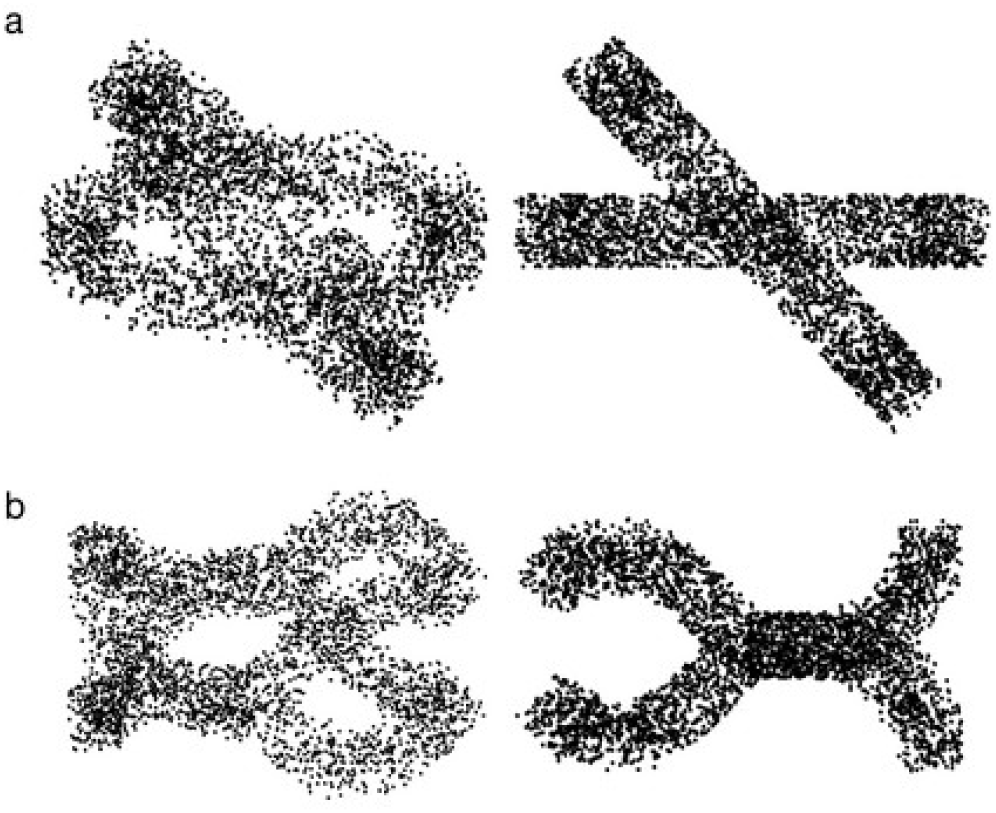
Stimuli. (a) Symmetric rings were presented during spontaneous reversals. (b) Asymmetric rings were presented during replayed reversals.

Differential activation z-scores in Fig.8(C,E,G) are calculated from the differential dPSTH z-scores. Fig.8(C) shows a further z-score calculated from the product *dX*(*R, τ*) = *dX*_*z*_(*A, τ*)*R*_nrm_(*A*) using the raw signal *dX*_*z*_(*A, τ*) described above. To calculate the further z-score *R*_nrm_(*A*) is shuffled across areas *A* and the surrogates *dX*(*R, τ*)_*surr*_ obtained. This is performed 1000 times to produce 1000 surrogate values and the z-score value calculated. Similarly Fig.8(E) shows a z-score calculated from the product *dP*_1*~*5_(*R, τ*) = *dP*_*z*,1*~*5_(*A, τ*)*R*_nrm_(*A*) using the dPSTH_*z*_ from the sum over the first five PCs, *dP*_*z*,1*~*5_(*A, τ*) described above and Fig.8(G) shows a z-score calculated from the product *dF*_1_(*R, τ*) = *dF*_*z*,1_(*A, τ*)*R*_nrm_(*A*). The correlation matrices shown in Fig.8(D,F,H) are the cross-correlations across *τ* between the networks *R* of the *dQ*(*R, τ*) for, respectively, the signal, *dX*(*R, τ*), the combination of the first five PCs, *dP*_1*~*5_(*R, τ*), and the first FC, *dF*_1_(*R, τ*).

## Declaration of the use of AI

No AI tools were used in planning or carrying out the research: stimulus generation, data acquisition, preprocessing, and all analyses and figures were produced with the software described in the Methods (Psychtoolbox-3, FSL, BrainWavelet, MRIcron, and custom MATLAB code) without AI assistance. In writing the paper, AI (or other tools) were used to check spelling and grammar; we used AI to reword our draft text; and we used AI to generate parts of the paper’s text, and verified its accuracy. Specifically, large language models were used to integrate, structure, and cross-reference multiple earlier manuscript versions, to check the internal consistency of the text before submission, and to copy-edit for clarity. AI played no role in generating figures. The exact AI tools were Anthropic’s Claude (Claude Opus 4.7, Claude Opus 5, Claude Opus 5.5, Claude Sonnet 5 and Claude Fable 5.1 models, accessed through Claude Code) and OpenAI’s Codex (gpt-5.6-sol and gpt-6-astra models, accessed through the Codex CLI). All scientific content, analyses, interpretations, citations, and final wording were authored, verified, and approved by the human authors, who take full responsibility for the content of the publication.

## Data and Code Availability

The fMRI dataset analysed in this study was acquired at the Centre for Neuroscientific Innovation and Technology (ZENIT), Magdeburg, under ethics permits 29/21 and 14/23 issued by the Ethics Commission of the Faculty of Medicine, Otto-von-Guericke University, Magdeburg (see Methods). De-identified single-subject time series at the MD758 parcellation level, perceptual-report event times, and the derived principal-component and noise-correlation matrices that underlie Figs. 1–8 will be deposited at the Open Science Framework (OSF) upon acceptance, with the repository URL and DOI provided at that time, for non-commercial research use, in accordance with the consent obtained from participants. Raw, individual-level fMRI volumes are not publicly shareable under the original consent terms but may be requested from the corresponding authors for bona fide research collaborations, subject to a data-use agreement and review by the Otto-von-Guericke University ethics committee. Analysis code (MATLAB scripts implementing the report-locked PSTH construction, principal component analysis, time-resolved noise-correlation analysis, and surrogate generation procedures described in the Methods) will be deposited in the same OSF repository. No new general-purpose code libraries were developed for this study: preprocessing relied on standard FSL (FMRIB, Oxford, UK), MRIcron, and BrainWavelet pipelines, with custom analysis performed in MATLAB (Mathworks) as specified in the Methods.

## Author Contributions

**Adam Ponzi:** Conceptualization, Data curation, Formal analysis, Methodology, Software, Visualization, Writing – original draft, Writing – review & editing. **Katsunori Miyahara:** Writing – original draft, Writing – review & editing. **Keisuke Suzuki:** Funding acquisition, Project administration, Supervision, Writing – review & editing.

## Acknowledgements

The authors are grateful to Stepan Aleshin, Ehsan Kakaei and Jochen Braun for supplying the data and useful discussions.

## Supplementary Material

**Figure S1:**
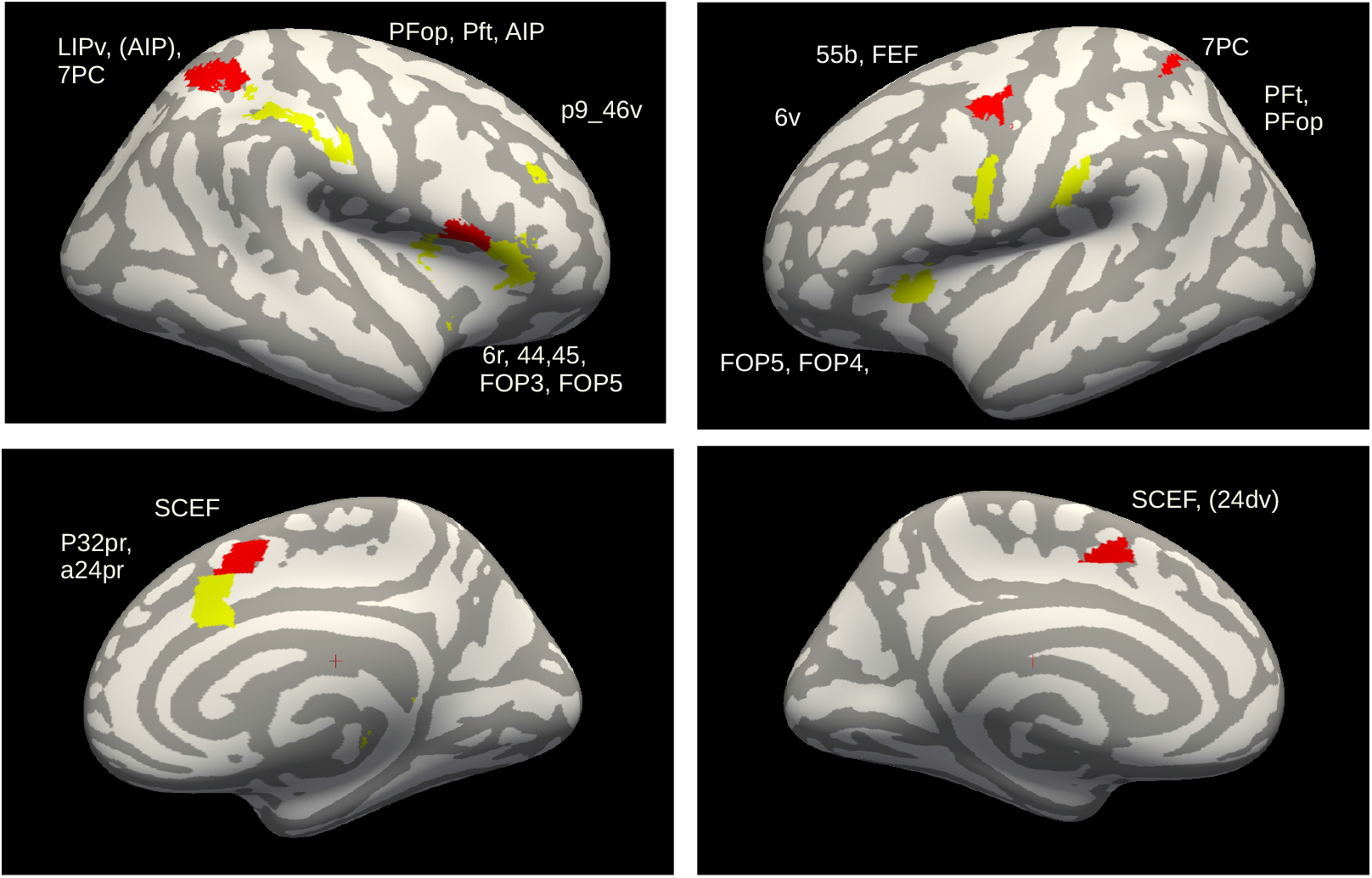
Areas significantly differentially activated in spontaneous compared to replayed perceptual switching using the raw signal, dSig, at *τ* = 3 s post-report, corresponding to Fig.2(A). Red (yellow) areas: actual dPSTH more positive than 3000 (2999) surrogates. Calculations use MD758 parcellation; areas labelled according to the HCP 360-area parcellation [38, 60].

**Figure S2:**
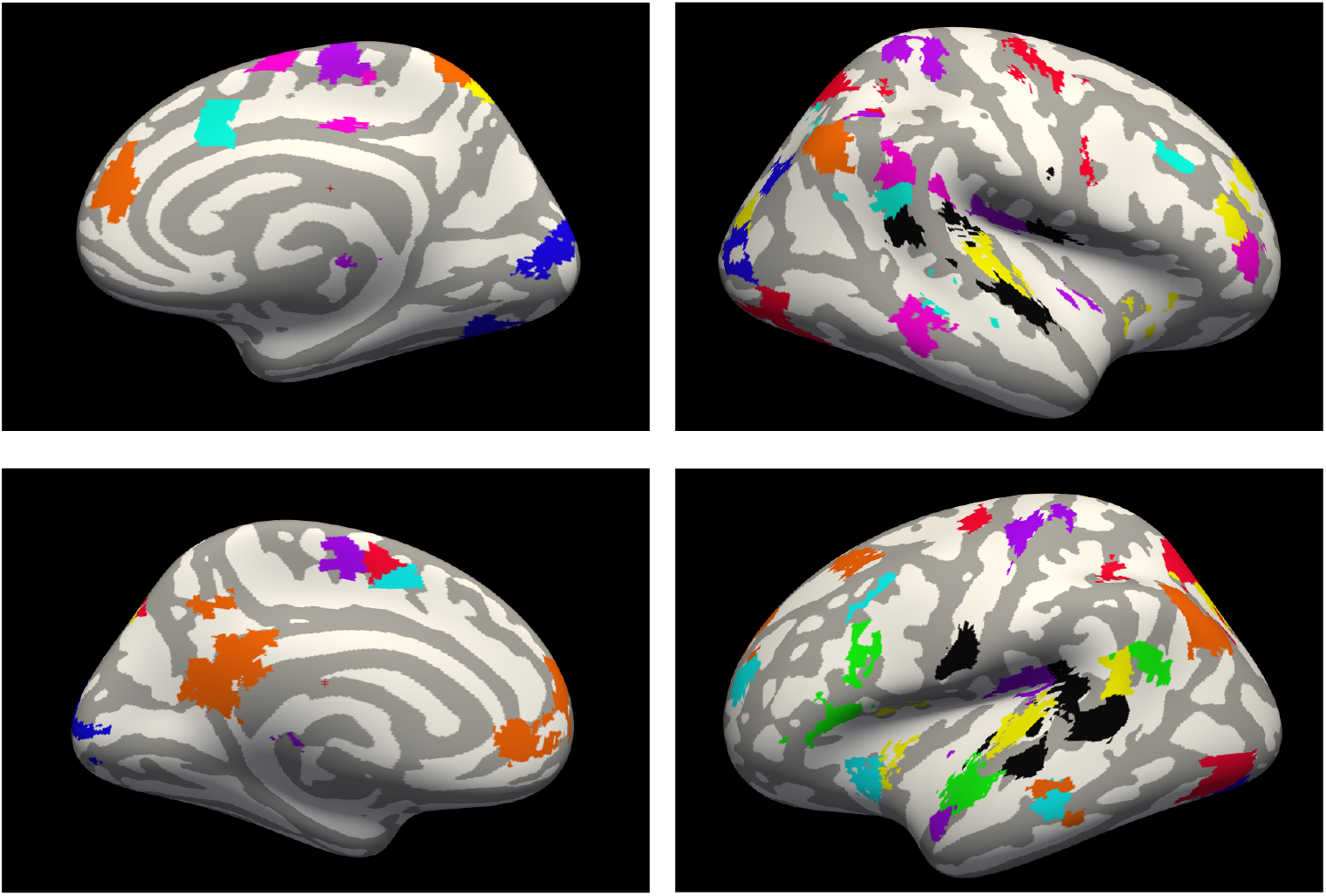
Resting-state networks (RSNs) used here, from [47]: (1) dorsal attention DAN (red), (2) ventral attention VAN (pink), (3) sensorimotor SMN (purple), (4) visual VIS (blue), (5) frontoparietal FPN (cyan), (6) language LAN (green), (7) auditory AUD (black), (8) default mode DMN (orange), (9) belonging to more than one network (yellow).

**Figure S3:**
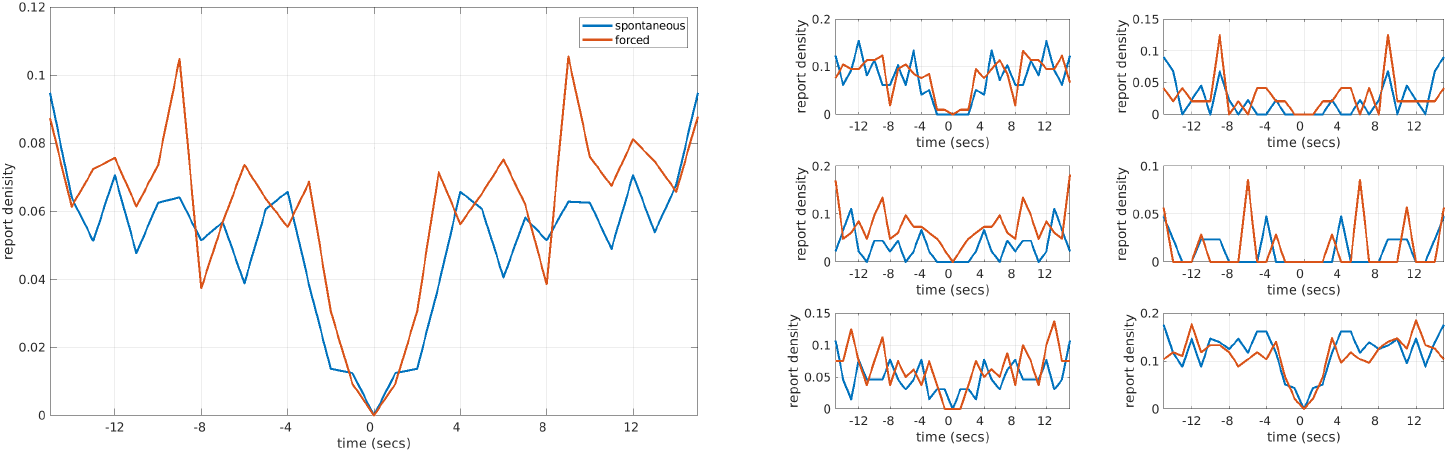
Perceptual report interval density given a report at *t* = 0 (the unity peak at *t* = 0 is removed for clarity). Left: average across subjects, spontaneous (blue) and replayed (orange; labelled “forced” in the legend as a legacy plotting label). Right: individual subjects. Spontaneous and replayed conditions show similar report intervals.

